# New routes for PTP1B allosteric inhibition by multitemperature crystallography, fragment screening and covalent tethering

**DOI:** 10.1101/218966

**Authors:** Daniel A. Keedy, Zachary B. Hill, Justin T. Biel, Emily Kang, T. Justin Rettenmaier, Jose Brandao-Neto, Nicholas M. Pearce, Frank von Delft, James A. Wells, James S. Fraser

**Affiliations:** Department of Bioengineering and Therapeutic Sciences, University of California, San Francisco, San Francisco, CA, USA; Department of Pharmaceutical Chemistry, University of California, San Francisco, San Francisco, CA, USA; Cellular and Molecular Pharmacology, University of California, San Francisco, San Francisco, CA, USA; present address: Jnana Therapeutics, Cambridge, MA, USA; Diamond Light Source, Harwell Science and Innovation Campus, Didcot OX11 0DE, England; Crystal & Structural Chemistry Group, Bijvoet Center for Biomolecular Research, Utrecht University, 3584 CH Utrecht; Structural Genomics Consortium (SGC), University of Oxford, Oxford OX3 7DQ, England; Department of Biochemistry, University of Johannesburg, Aukland Park, Johannesburg 2006, South Africa

**Keywords:** allostery, X-ray crystallography, conformational heterogeneity, ligand binding, phosphatase

## Abstract

Allostery is an inherent feature of proteins and provides alternative routes to regulating function. Small-molecule allosteric inhibitors are often desirable; however, it remains challenging to identify surface sites in proteins which can bind small molecules and modulate function. We identified new allosteric sites in protein tyrosine phosphatase 1B (PTP1B) by combining multiple-temperature X-ray crystallography experiments and structure determination from hundreds of individual small-molecule fragment soaks. New modeling approaches reveal “hidden” low-occupancy conformational states for protein and ligands. Our results converge on a new allosteric site that is conformationally coupled to the active-site WPD loop, a hotspot for fragment binding, not conserved in the closest homolog, and distinct from other recently reported allosteric sites in PTP1B. Targeting this site with covalently tethered molecules allosterically inhibits enzyme activity. Overall, this work demonstrates how the ensemble nature of macromolecular structure revealed by multitemperature crystallography can be exploited for developing allosteric modulators.

## Introduction

The most common way to inhibit the activity of a protein is to target its active site with a small molecule. Unfortunately, because active sites are often conserved among homologous proteins, many active-site inhibitors bind non-specifically, leading to off-target cellular effects [1]. Moreover, active sites are often highly polar, so their complementarily polar inhibitors often suffer from poor bioavailability [2]. These problems are exemplified by the challenge of identifying inhibitors for protein tyrosine phosphatases [3]. An allosteric inhibitor that binds to a less-conserved and less-polar surface site could bypass these limitations. This prospect is appealing because allostery may be inherent to nearly all protein structures [4,5]. Despite the promise of allosteric inhibition, it remains challenging to pinpoint sites on the extensive and often flexible surface of a protein structure that are both allosterically coupled to the active site and bindable by small molecules.

To understand the relationship between small molecule binding and allosteric coupling to the active site, we examined the archetypal protein tyrosine phosphatase, PTP1B, as a model system. PTP1B is highly validated as a therapeutic target for diabetes [6] and cancer [7], and has also been linked to Rett syndrome [8]. Several approaches have been used to selectively target the active site of PTP1B, such as linking non-hydrolyzable phosphotyrosine (pTyr) analogs that bind the active site with small-molecule fragments that bind in nearby, less conserved sites [9]. Unfortunately, to date, no active-site inhibitors have reached the clinical stage, leading some to label PTP1B “undruggable”. Two classes of compounds have been identified that allosterically inhibit PTP1B. The first class of compounds are based on a benzbromarone (BB) scaffold and inhibit allosterically by binding to the space normally occupied by the regulatory C-terminal α7 helix [10]. Recent work combining mutagenesis, X-ray crystallography, NMR spectroscopy, and molecular dynamics simulations revealed how rotations of helix α3 and a discrete switch of the catalytic WPD loop are impacted by these BB allosteric inhibitors [11]. The second class are natural products that bind to multiple sites that primarily involve the disordered C-terminus [7]. Despite these discoveries, the molecules suffer from limitations that prevent their translation to the clinic. Therefore, there remains a need for more general methods for identifying small-molecule binding sites that can allosterically modulate PTP1B or other “undruggable” systems.

We have addressed the challenge of discovering new allosteric sites and small-molecule inhibitors for PTP1B by taking advantage of two new techniques in X-ray crystallography that reveal minor conformational states of protein and ligands. First, multitemperature crystallography [12] can reveal previously hidden alternative conformations that enable biological functions. Here we use this approach in PTP1B to reveal coupled conformations that colocalize in a concerted allosteric network. This approach validates many aspects of the recently characterized allosteric network that responds to BB inhibitors [11] and identifies new, allosterically coupled binding sites. Second, high-throughput small-molecule fragment soaking and structure determination [13] has enabled new algorithms for revealing low-occupancy ligands [14]. We use this approach to comprehensively canvas the PTP1B surface with 1,627 small-molecule fragments, 110 of which were resolved by crystallography. This independent approach converges on the same promising allosteric sites as multitemperature crystallography. Finally, we interrogate one of these new allosteric sites with covalently tethered small molecules [15] to confirm a functional link to enzyme activity. Overall, our work opens new avenues for developing potent allosteric inhibitors for PTP1B, and provides a more general roadmap for characterizing and rationally exploiting coupled conformational heterogeneity to control protein function.

## Results

### Multitemperature crystallography to identify allosterically coupled residues

To identify new allosteric sites in PTP1B that may communicate with the active site, we searched for regions of the protein whose conformational heterogeneity is coupled to that of the active site. We began by examining the conformational heterogeneity of the active-site WPD loop; transition of this loop from the open to the closed state is rate-limiting for catalysis [16]. In the only available apo crystal structure of PTP1B in which the WPD loop is free from crystal-lattice contacts (PDB ID 1sug) [17], the loop is modeled in the closed state. However, low-contour electron density can reveal hidden alternative conformations in protein crystal structures [18,19]. We therefore investigated the electron density near the WPD loop in the apo structure more closely (**Figure 1B**).

**Figure 1:**
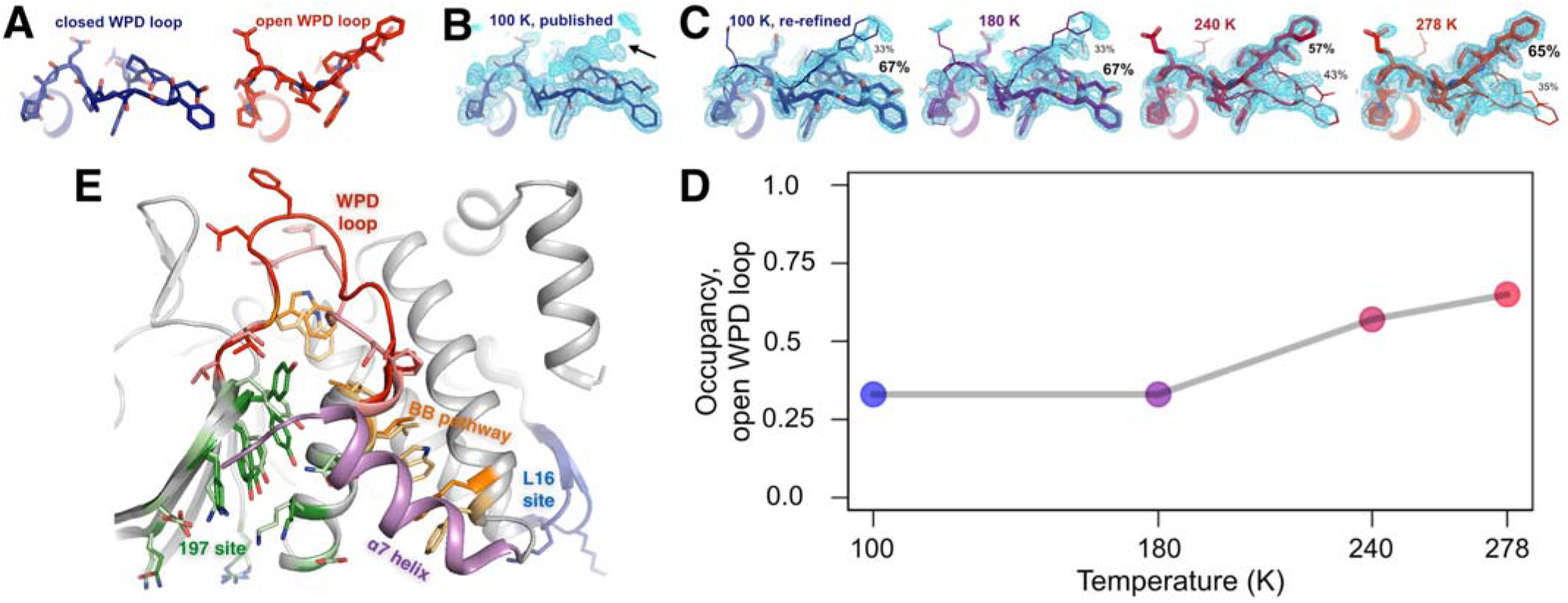
The conformational ensemble of the active-site WPD loop and allosterically coupled regions. **(A)** The active-site WPD loop in PTP1B adopts either a closed conformation (example from PDB ID 1sug) or an open conformation (example from PDB ID 1t49). View from the “front side” of PTP1B. **(B)** In the previously published apo structure of PTP1B, solved at 100 K (PDB ID 1sug), 0.8 σ 2Fo-Fc electron density (cyan) supports the modeled closed conformation, but substantial electron density remains unexplained (arrow). **(C)** Adding the open conformation of the WPD loop as a secondary conformation at partial occupancy accounts for this electron density. In structures solved at different elevated temperatures, electron density for the open conformation becomes more prominent as its occupancy (labeled) relative to the closed conformation increases. **(D)** The occupancy of the open conformation increases non-linearly with temperature. **(E)** Overall roadmap of allostery on the “back side” of PTP1B, with the new 197 allosteric site and loop 16 (L16) allosteric site highlighted in the context of the larger allosteric network including the previously established BB pathway, a7 helix, and WPD loop. Sidechains are shown in stick representation for several key residues in the WPD loop and allosteric regions. For those residues with alternative conformations at 278 K, both open-state (darker hues) and closed-state (lighter hues) conformations are shown.

### Figure 1 - Movie S1

Movie version with five scenes: **Figure 1B** then **Figure 1C**.

### Figure 1 - Movie S2

Movie version of **Figure 1E**.

Surprisingly, upon closer inspection, the electron density strongly suggests a significant population for the open state as well (**Figure 1C, left**). Our re-refined model with both open and closed states as alternative conformations visually accounts for the electron density around this loop much better than the original model (**Figure 1C, left** and **Figure 1 - Figure S2**). By contrast, when we re-refined 36 other available crystal structures of PTP1B complexed with active-site inhibitors using both open and closed loop states as putative alternative conformations, Fo-Fc difference electron density and the bimodal distribution of refined occupancies indicated the single-state models were a better fit (**Figure 1 - Figure S1**). These results suggest that, even in the crystal, apo PTP1B samples both WPD loop states and that active-site inhibitors then lock the loop either fully open or fully closed.

**Figure 1 - Figure S1:**
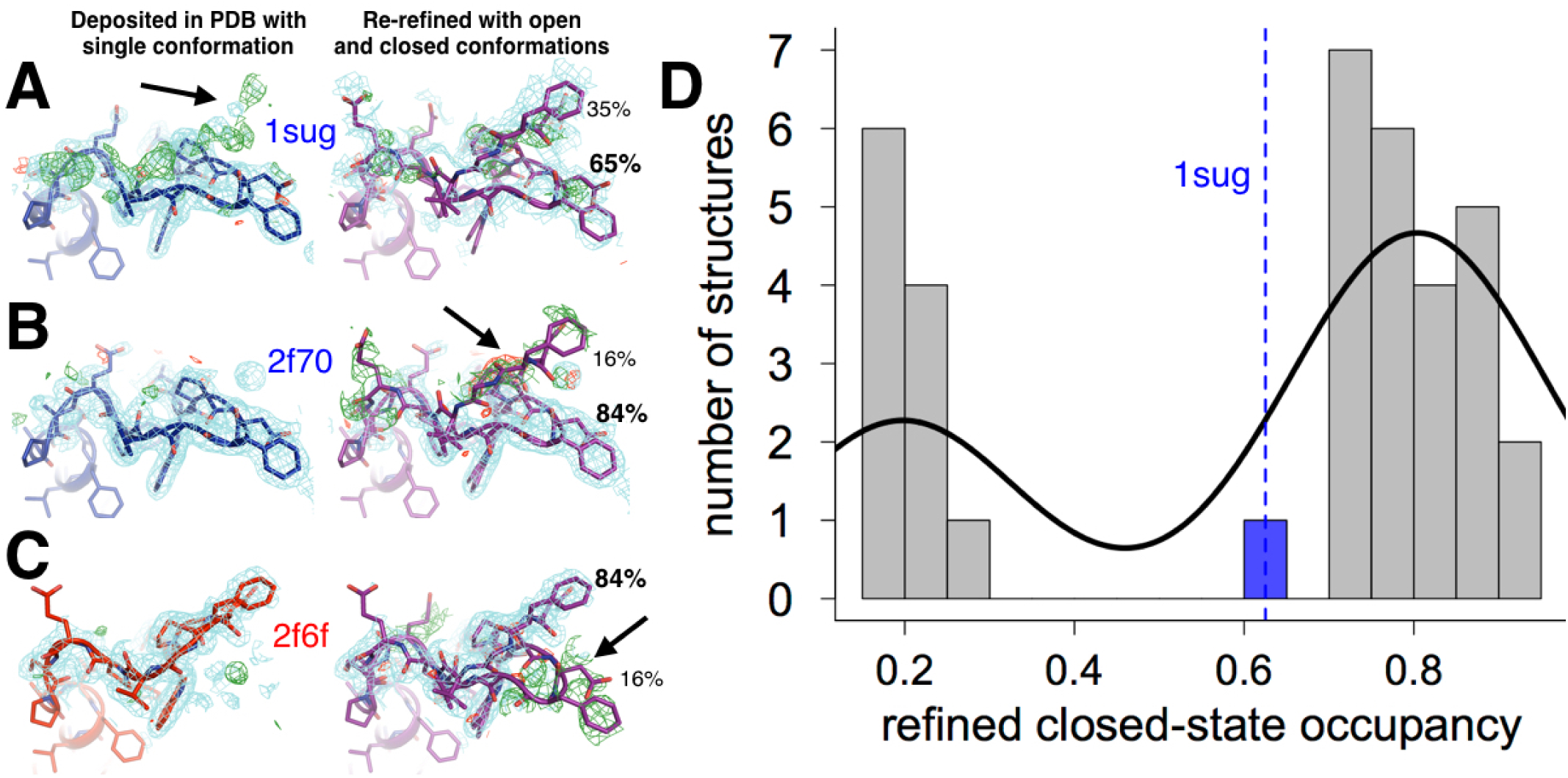
The WPD loop adopts multiple conformations only in the absence of inhibitors. 36 structures from the PDB in the same space group as our datasets (P3_12_1), originally modeled as only-open or only-closed, were naively re-refined with open and closed WPD loops modeled as alternative conformations with occupancies constrained to sum to 1. Almost all models have an active-site inhibitor bound. In P3_1_21 the loop is free from any crystal contacts. **(A-C)** Representative examples of structures with the WPD loop originally modeled as closed or open (left, blue or red), and our re-refined dual-conformation models (right, purple). 1.0 σ 2Fo-Fc (cyan) and +/- 3.0 σ Fo-Fc electron density (green/red) are shown. **(A)** PDB ID 1sug: apo PTP1B; originally closed. 2Fo-Fc electron density supports the dual-conformation model; residual Fo-Fc density reflects partial-occupancy waters (**Figure 1 - Figure S2**). **(B)** PDB ID 2f70: PTP1B in complex with an active-site inhibitor; originally closed. Fo-Fc electron density suggests the dual-conformation model is a poor fit. **(C)** PDB ID 2f6f: PTP1B in complex with another active-site inhibitor; originally open. Fo-Fc electron density suggests the dual-conformation model is a poor fit. **(D)** Loop occupancies for the re-refined dual-conformation models are bimodal: either ~fully open (left) or ~fully closed (right). The curve is a fit to the histogram using kernel density estimation, and is shown to emphasize the bimodal nature of the distribution. The only exception is the one published apo structure in this space group (1sug), which can be successfully refined with partial open and closed occupancies.

**Figure 1 - Figure S2:**
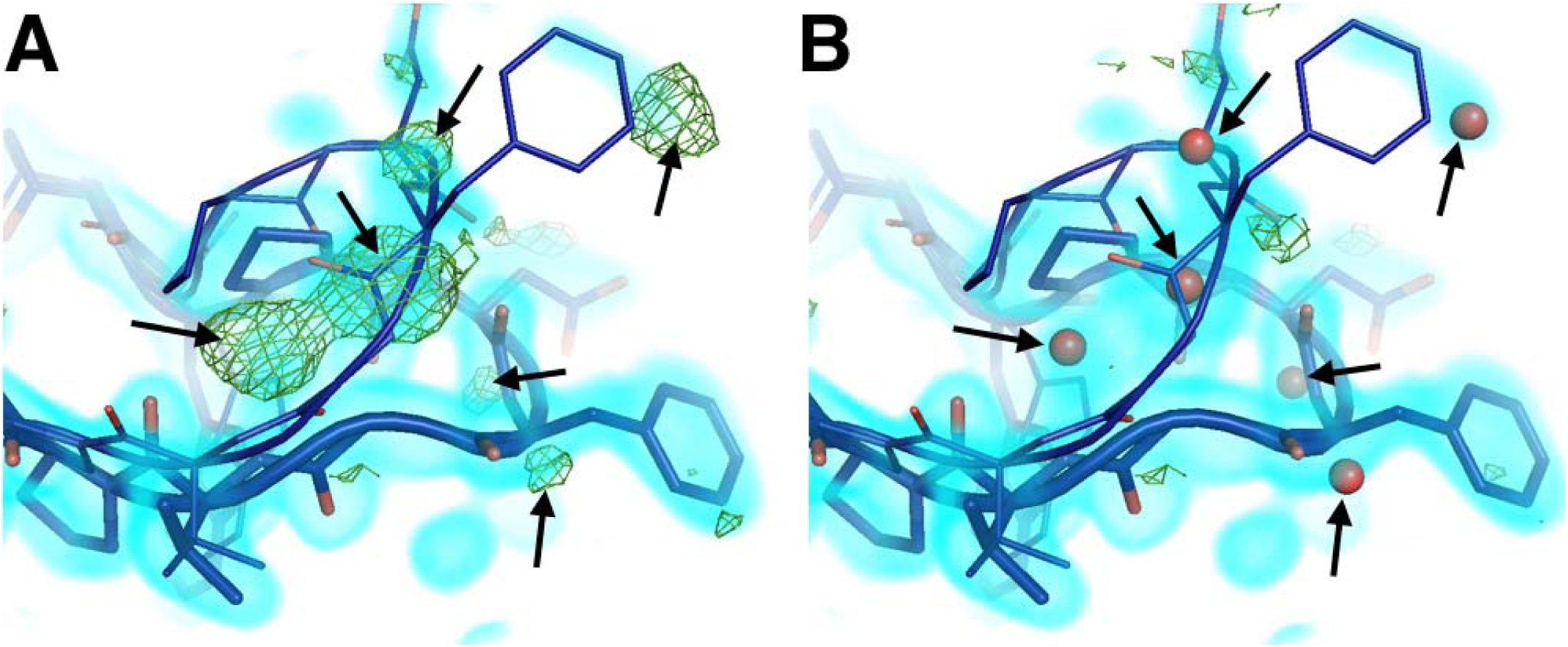
Mutually exclusive partial-occupancy protein and solvent atoms complicate model building. **(A)** Although both closed and open WPD loop states are supported by 0.8 σ 2Fo-Fc electron density (cyan volume) in the 100 K apo PTP1B structure, several +3.2 σ Fo-Fc electron density (green mesh) remain unexplained (arrows). **(B)** Partial-occupancy waters (red spheres) with the opposite “altloc” alternative conformation label as the closest protein atoms (“A” vs. “B”) dramatically reduce the difference density and better explain the data. See [17]. The waters modeled here resemble water networks from either the closed-only (2f70) or the open-only (2f6f) structures.

To better characterize the conformational heterogeneity of the WPD loop in apo PTP1B, we collected X-ray datasets at several elevated temperatures (180, 240, and 278 K) in addition to the cryogenic 100 K model from the PDB, all at better than 2 resolution (**Table 1**). Each complete dataset was obtained from a single crystal, and crystallographic statistics indicated that radiation damage was not a concern even at the elevated temperatures [20] (**Table 1 - Figure S1**). We built an initial multiconformer model for each temperature using the automated algorithm qFit [21]. These models are parsimonious in that each atom has alternative positions only if justified by the experimental data, and a single position otherwise. We then manually refined alternative conformations for protein, buffer components, and solvent. In particular, we took advantage of the wealth of available structures of PTP1B in the PDB [22] to sample coordinates for putative alternative conformations; in many cases, these conformations explained missing regions with positive Fo-Fc electron density that would have otherwise been difficult to model.

**Table 1:** Crystallographic statistics for multitemperature and mutant X-ray datasets. Overall statistics given first (statistics for highest-resolution bin in parentheses). WT apo, 100 K (*): statistics are taken from our remodeled structure where appropriate, or from the original PDB ID 1sug when possible otherwise, or given as “—” where unavailable.

|  | WT apo,<br>100 K (*) | WT apo,<br>180 K | WT apo,<br>240 K | WT apo,<br>278 K | WT with<br>BB3, 278<br>K | K197C<br>apo, 100 K | K197C<br>tethered to<br>2, 100 K |
| --- | --- | --- | --- | --- | --- | --- | --- |
| PDB ID | 6B90 | 6B8E | 6B8T | 6B8X | 6B8Z | 6BAI | 6B95 |
| Number of crystals used | 1 | 1 | 1 | 1 | 1 | 1 | 1 |
| Wavelength (Å) | 0.8110 | 1.115869 | 1.115869 | 0.9795 | 1.11583 | 1.11583 | 1.11583 |
| Resolution range (Å) | 33.60 - 1.95 (2.02 - 1.95) | 19.18 - 1.82 (1.89 - 1.82) | 62.54 - 1.85 (1.92 - 1.85) | 31.31 - 1.74 (1.80 - 1.74) | 43.88 - 1.8 (1.86 - 1.8) | 75.57 - 1.95 (2.02 - 1.95) | 43.98 - 1.95 (2.02 - 1.95) |
| Space group | P 31 2 1 | P 31 2 1 | P 31 2 1 | P 31 2 1 | P 31 2 1 | P 31 2 1 | P 31 2 1 |
| Unit cell (Å, °) | 88.12<br>88.12<br>103.90 90<br>90 120 | 88.57<br>88.57<br>104.32 90<br>90 120 | 89.44<br>89.44<br>106.00<br>90 90<br>120 | 89.52<br>89.52<br>106.25 90<br>90 120 | 89.65<br>89.65<br>106.39 90<br>90 120 | 87.27<br>87.27<br>104.10 90<br>90 120 | 87.96 87.96<br>104.63 90<br>90 120 |
| Total reflections | --- | 339461<br>(33574) | 256701<br>(25484) | 299041<br>(30678) | 771858<br>(76158) | 653798<br>(63759) | 170094<br>(16845) |
| Unique reflections | 34486<br>(3371) | 42866<br>(4223) | 42343<br>(4156) | 50486<br>(4960) | 46311<br>(4561) | 33409<br>(3296) | 34596<br>(3387) |
| Multiplicity | 7.4 (---) | 7.9 (7.9) | 6.1 (6.1) | 5.9 (6.2) | 16.7 (16.7) | 19.6 (19.3) | 4.9 (5.0) |
| Completeness (%) | 99.9<br>(100.0) | 99.9<br>(100.0) | 100.0<br>(100.0) | 99.0<br>(98.3) | 100.0<br>(100.0) | 98.06<br>(98.51) | 99.81<br>(99.88) |
| Mean I/sigma(I) | 26.4<br>(4.4) | 21.40<br>(1.46) | 15.89<br>(1.46) | 17.15<br>(1.49) | 11.69<br>(0.75) | 25.06<br>(0.94) | 12.04<br>(0.81) |
| Wilson B (Å<sup>2</sup>) | 24.12 | 23.69 | 28.48 | 19.48 | 34.96 | 46.3 | 38.82 |
| R<sub>merge</sub> | 0.073<br>(0.515) | 0.089<br>(1.483) | 0.079<br>(1.215) | 0.098<br>(1.209) | 0.148<br>(2.664) | 0.076<br>(3.056) | 0.088<br>(1.852) |
| R<sub>meas</sub> | --- | 0.096<br>(1.585) | 0.086<br>(1.329) | 0.108<br>(1.318) | 0.153<br>(2.748) | 0.078<br>(3.138) | 0.098<br>(2.071) |
| $R_{\text{pim}}$ | --- | 0.034<br>(0.556) | 0.035<br>(0.535) | 0.044<br>(0.520) | 0.038<br>(0.672) | 0.018<br>(0.705) | 0.044<br>(0.916) |
| CC1/2 | --- | 0.999<br>(0.520) | 0.999<br>(0.560) | 0.999<br>(0.496) | 0.997<br>(0.366) | 1.000<br>(0.438) | 0.999<br>(0.292) |
| CC* | --- | 1.000<br>(0.827) | 1.000<br>(0.847) | 1.000<br>(0.814) | 0.999<br>(0.732) | 1.000<br>(0.781) | 1.000<br>(0.672) |
| Reflections used<br>in refinement | 34486<br>(3371) | 42862<br>(4223) | 42343<br>(4156) | 50486<br>(4959) | 46309<br>(4561) | 33302<br>(3296) | 34576<br>(3386) |
| Reflections used<br>for $R_{\text{free}}$ | 1356<br>(128) | 1689<br>(165) | 1669<br>(163) | 1993<br>(196) | 1820<br>(176) | 1310<br>(135) | 1362<br>(135) |
| $R_{\text{work}}$ | 0.1580<br>(0.1881) | 0.1708<br>(0.2781) | 0.1674<br>(0.2760) | 0.1752<br>(0.3511) | 0.1675<br>(0.3116) | 0.2061<br>(0.3365) | 0.1858<br>(0.3295) |
| $R_{\text{free}}$ | 0.1926<br>(0.2090) | 0.1997<br>(0.2911) | 0.2123<br>(0.2869) | 0.2059<br>(0.3888) | 0.1978<br>(0.3017) | 0.2569<br>(0.3292) | 0.2307<br>(0.3686) |
| CC<sub>work</sub> | --- | 0.963<br>(0.753) | 0.958<br>(0.789) | 0.964<br>(0.673) | 0.964<br>(0.709) | 0.934<br>(0.618) | 0.957<br>(0.574) |
| CC<sub>free</sub> | --- | 0.948<br>(0.744) | 0.908<br>(0.677) | 0.948<br>(0.739) | 0.961<br>(0.757) | 0.994<br>(0.484) | 0.938<br>(0.294) |
| Non-H atoms | 2997 | 3021 | 3149 | 3260 | 2709 | 2502 | 2541 |
| Macromolecule<br>atoms | 2657 | 2687 | 2875 | 3011 | 2460 | 2377 | 2370 |
| Ligand atoms | 46 | 40 | 34 | 24 | 82 | 16 | 36 |
| Solvent atoms | 294 | 294 | 240 | 225 | 167 | 109 | 135 |
| Protein residues | 298 | 298 | 299 | 298 | 289 | 282 | 285 |
| RMS bonds ($\text{\AA}$) | 0.010 | 0.011 | 0.011 | 0.012 | 0.011 | 0.015 | 0.013 |
| RMS angles ($^{\circ}$) | 1.01 | 1.02 | 1.10 | 1.08 | 1.14 | 1.30 | 1.15 |
| Ramachandran favored (%) | 97.64 | 97.64 | 97.98 | 95.95 | 97.19 | 95.00 | 95.76 |
| Ramachandran allowed (%) | 2.03 | 2.03 | 1.68 | 3.72 | 2.11 | 4.64 | 2.83 |
| Ramachandran outliers (%) | 0.34 | 0.34 | 0.34 | 0.34 | 0.70 | 0.36 | 1.41 |
| Rotamer outliers (%) | 1.35 | 2.67 | 1.87 | 2.38 | 2.19 | 6.44 | 4.91 |
| Clashscore | 2.79 | 3.31 | 3.62 | 2.98 | 2.79 | 6.06 | 3.36 |
| MolProbity score | 1.24 | 1.52 | 1.36 | 1.65 | 1.47 | 2.29 | 1.95 |
| Average B ($\text{\AA}^2$) | 29.35 | 30.22 | 36.50 | 28.06 | 43.58 | 55.81 | 49.66 |
| Average B, macromolecule ($\text{\AA}^2$) | 28.14 | 29.11 | 35.51 | 27.01 | 43.30 | 56.02 | 49.53 |
| Average B, ligands ($\text{\AA}^2$) | 47.43 | 48.08 | 60.29 | 67.33 | 41.13 | 45.41 | 53.91 |
| Average B, solvent ($\text{\AA}^2$) | 37.46 | 37.90 | 44.96 | 38.01 | 48.96 | 52.76 | 50.82 |

**Table 1 - Figure S1:**
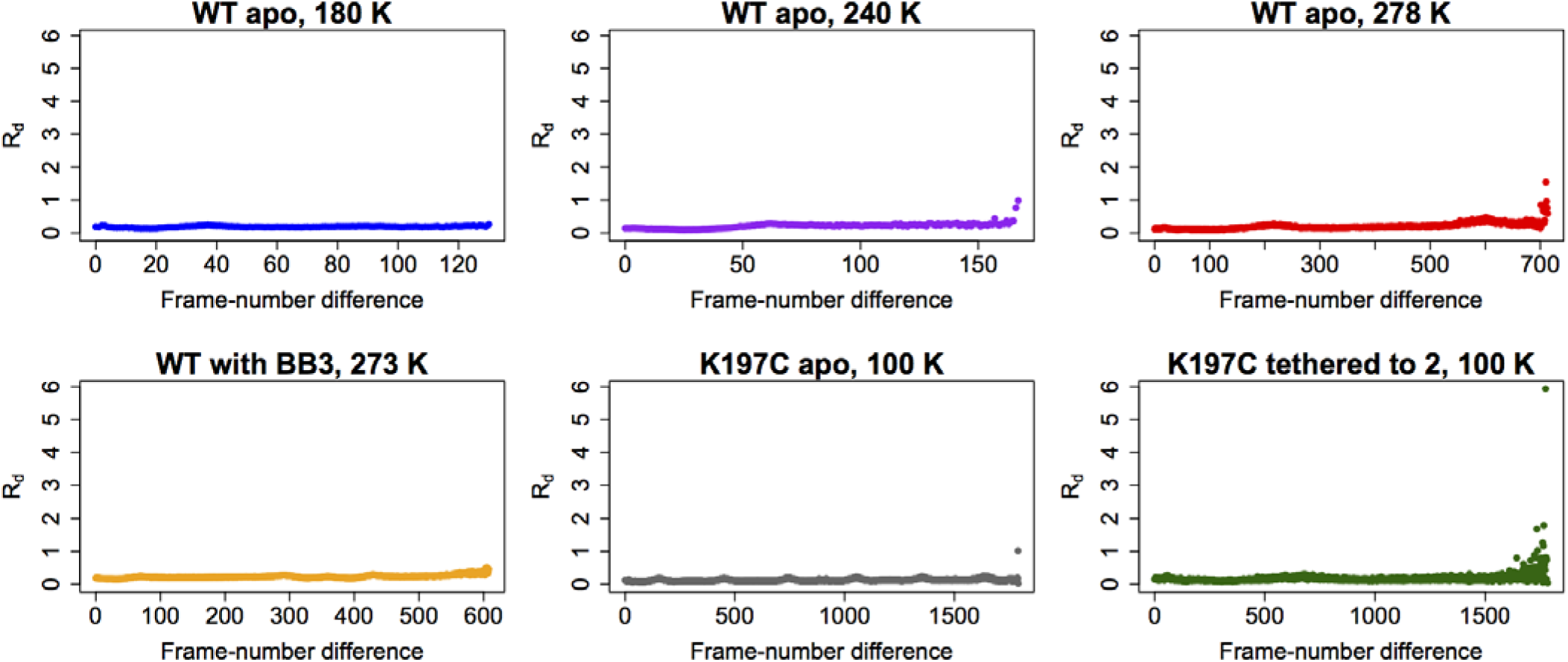
Radiation damage is minimal across all datasets. Plots of R_d_ [20] versus frame-number difference for each dataset reveal little evidence for radiation damage. Statistics for 100 K, which is from PDB ID 1sug, are not shown.

The WPD loop adopts both the open and closed conformations across this range (**Figure 1C**) and the the population of the open vs. closed states was sensitive to temperature (**Figure 1D**). The loop is approximately 67% closed at 100 K, but 65% open at 278 K. These occupancies evolve non-linearly [12] at intermediate temperatures.

Overall, we also observed temperature-dependent conformational heterogeneity for several other regions of PTP1B, including the previously characterized BB allosteric site plus a new “197 site” and “loop 16 16 (L16) site”. These regions are all contiguous in the structure (**Figure 1E**), suggesting that they together constitute an expanded collective allosteric network in PTP1B that is coupled to the WPD loop. The manner in which they are connected is described in detail in the following sections.

### Multitemperature crystallography of the BB allosteric site

To connect these multitemperature structures to known allosteric regulatory mechanisms, we first turned to a benzbromarone derivative compound (here referred to as BB2) that binds to an allosteric site >12 □ away from the active site and inhibits enzyme activity [10]. The authors described a series of induced conformational changes that begins with BB2 directly displacing Trp291 to disorder the entire C-terminal α7 helix, and ends with Phe191 χ2 dihedral-angle rotations clashing with the WPD loop anchor residue Trp179 to stabilize the open state. We tested the hypothesis that these allosterically inhibited conformations pre-exist in apo PTP1B by examining these regions in our multitemperature apo crystal structures. Indeed, in apo PTP1B the α7 helix is more ordered at lower temperatures but more disordered at higher temperatures (**Figure 2A**). Also, Trp179 and Phe191 adopt dual conformations at higher temperatures (**Figure 2B**) that coincide remarkably well with the apo and allosterically inhibited conformations (**Figure 2C**). We also see see alternative conformations at high temperatures for several residues within and directly flanking the WPD loop (Arg221, Pro185, Trp179, Phe269) which have been implicated as being important for a CH/π switch during WPD loop opening/closing [11] (**Figure 2 - Figure S1**). Multiple conformations for Leu192 were more difficult to detect at higher temperatures in apo PTP1B. This is likely because Leu192 shifts more subtly between the cryogenic apo and allosterically inhibited conformations, which is also consistent with a recent report that Leu192 is a relatively static inter-helical “wedge” [11]. Taken together, these results suggest that BB2 stabilizes a subset of pre-existing conformations in apo PTP1B.

**Figure 2:**
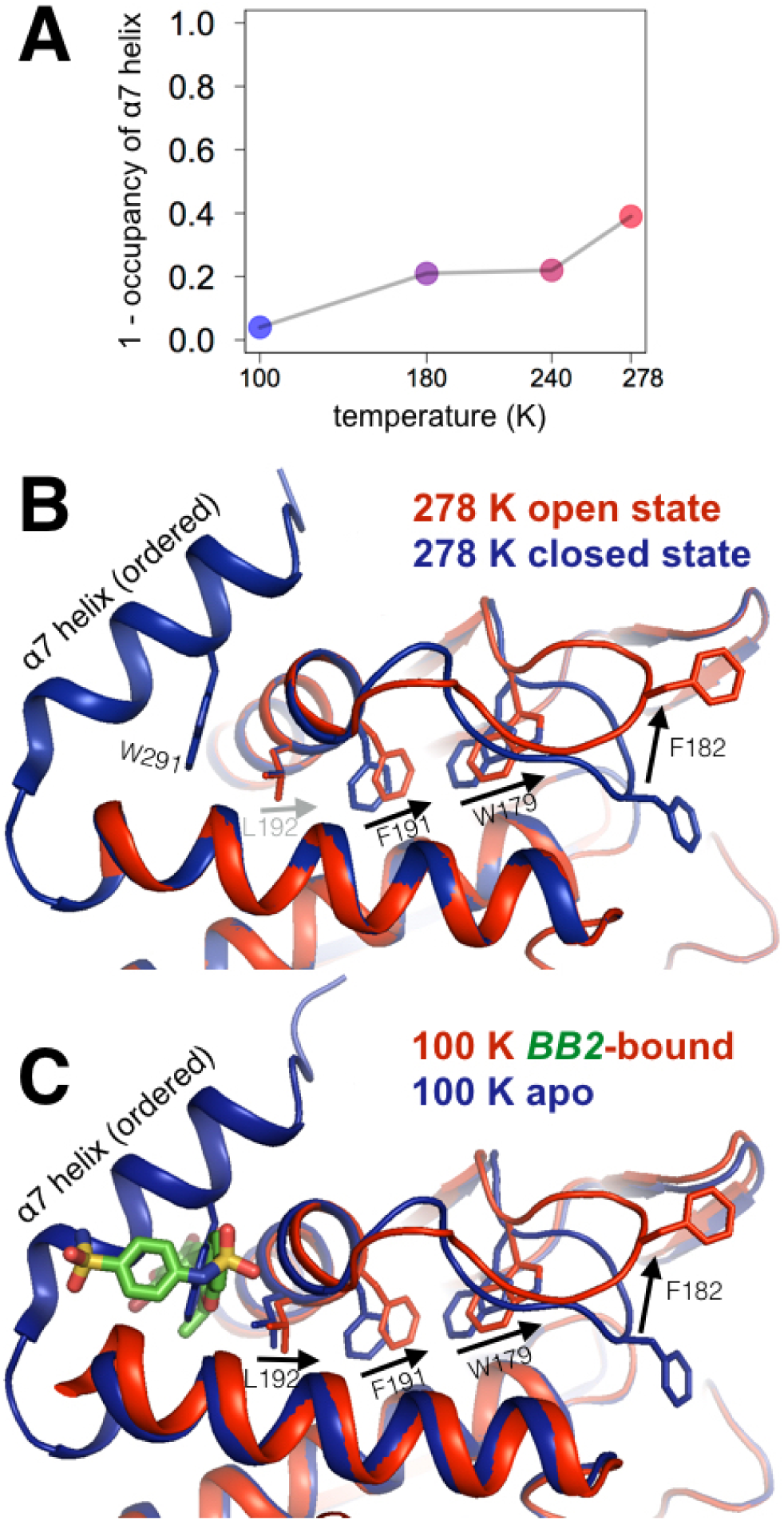
Multitemperature crystallography of apo PTP1B recapitulates an allosteric mechanism. **(A)** In apo PTP1B, the occupancy of the α7 helix decreases (i.e. the helix becomes more disordered) with temperature. The helix was modeled with one conformation and its occupancy was refined; the remaining occupancy corresponds to the disordered state of the helix. **(B)** Several residues that allosterically link α7 and the active-site WPD loop also undergo shifts with temperature. **(C)** These additional conformations match the state trapped by the allosteric inhibitor BB2 (PDB 1t49) [10] which binds >12 away from the active site.

**Figure 2 - Figure S1:**
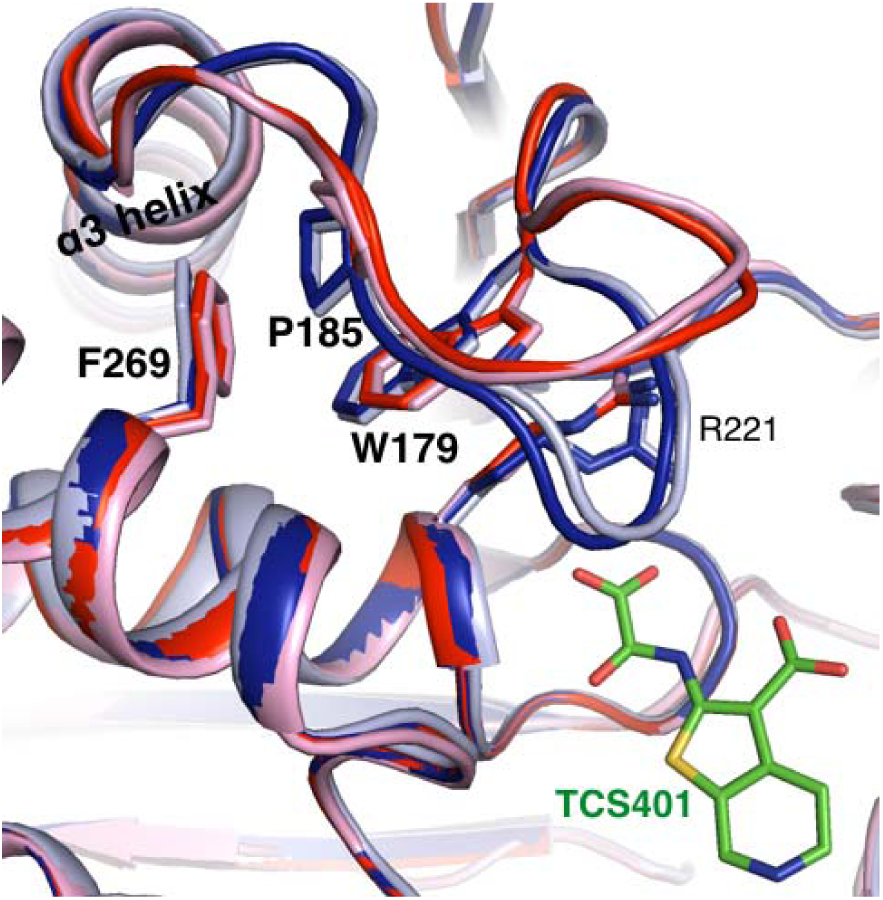
Alternative conformations in apo PTP1B recapitulate a reported conformational switching mechanism in the active site during WPD loop closing. Alternative conformations in our 278 K apo structure corresponding to the open state (red) vs. closed state (dark blue) recapitulate conformational changes for several residues (labeled) in the active site that undergo a CH/π-mediated switch between the open state (pink, 5k9v) and TCS401-inhibitor-bound closed state (light blue, 5k9w) [11].

We additionally solved a high-resolution (1.80 □, **Table 1**) room-temperature structure of PTP1B in complex with BB3 (which differs from BB2 only by an extra terminal aminothiazole group) and found it to be very similar to the cryogenic structures with BB3 (PDB ID 1t4j) and with BB2 (PDB ID 1t49) despite the difference in temperature (**Figure 2 - Figure S2**). However, two interesting features are evident at room temperature. First, at room temperature but not at cryogenic temperature, modeling BB3 with a single conformer leads to Fo-Fc difference electron density peaks at both ends of the molecule (**Figure 2 - Figure S3A**). To account for these peaks in the model, it is necessary to add a second alternative conformer, which includes a translation at one end and dihedral-angle changes at the other end (**Figure 2 - Figure S3B**). Chemical changes to BB3 designed to eliminate this remaining heterogeneity could potentially improve affinity and inhibition.

**Figure 2 - Figure S2:**
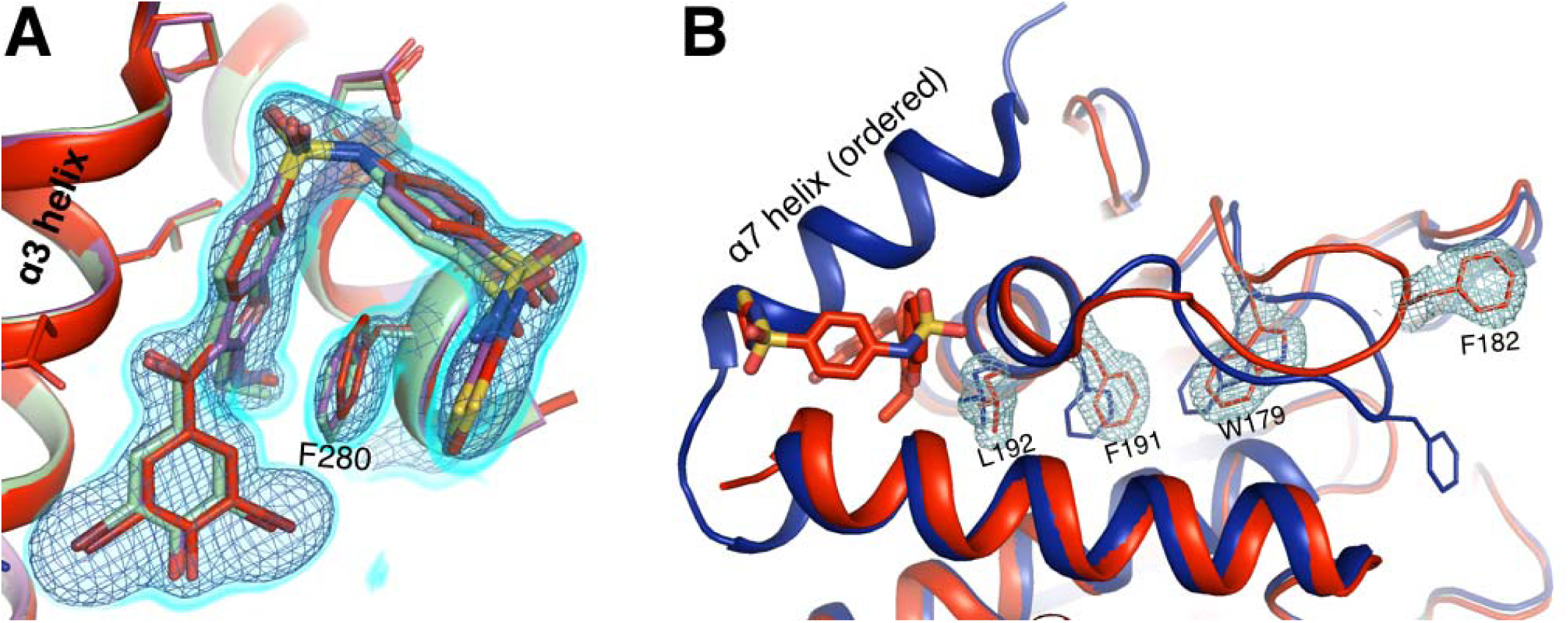
Allosteric inhibitor binding quenches conformational heterogeneity regardless of temperature. **(A)** The pose of BB3 in our new 1.80 □ room-temperature (273 K) structure (red) is well-justified by 2Fo-Fc electron density contoured at 0.75 σ (cyan volume) and at 1.5 σ (blue mesh). The pose is also essentially identical to the poses in previously published cryogenic structures with BB3 (1t4j, magenta) and BB2 (1t49, pale green), which is very similar in structure and inhibition to BB3 [10]. **(B)** 2Fo-Fc electron density contoured at 0.75 σ (cyan) supports only a single conformation along the previously characterized allosteric pathway, rather than the multiple conformations seen at 278 K from our multitemperature apo PTP1B series (**Figure BB_PATH**); a cryogenic structure in the closed state (1sug, blue) is shown for comparison. This confirms that inhibitor binding quenches conformational heterogeneity essentially completely, locking the protein in the open state.

Second, at room temperature we observe significant electron density just above BB3 (**Figure 2 - Figure S3C**). Modeling a reordered, non-helical conformation of α7 explains this density well, and places Trp291 in good position for aromatic stacking interactions with BB3 and other interactions with nearby sidechains on the α3 helix (**Figure 2 - Figure S3D**). Trp291 is displaced by BB3 or BB2 binding in a striking example of molecular mimicry [10] (**Figure 2C**). Our room-temperature data suggest that a subsequent reordering of the α7 polypeptide occurs, which may contribute to the affinity of BB3 for PTP1B. In contrast to our room-temperature data, electron density in this region is weak in the cryogenic structures with BB3 and BB2. However, in the cryogenic structure with BB1, a different derivative of the BB scaffold, α7 also reorders – but adopts a significantly different conformation than we observe at room temperature with BB3 (**S_BB3_RT_NOVEL_E,F**). Together, these results suggest that in addition to being a major allosteric hub when ordered [11], α7 is also quite malleable when disordered, and may interact in diverse ways with bound ligands – behavior which is similar to the mechanism proposed for inhibitors that bind via the disordered C-terminus beyond α7 [7].

**Figure 2 - Figure S3:**
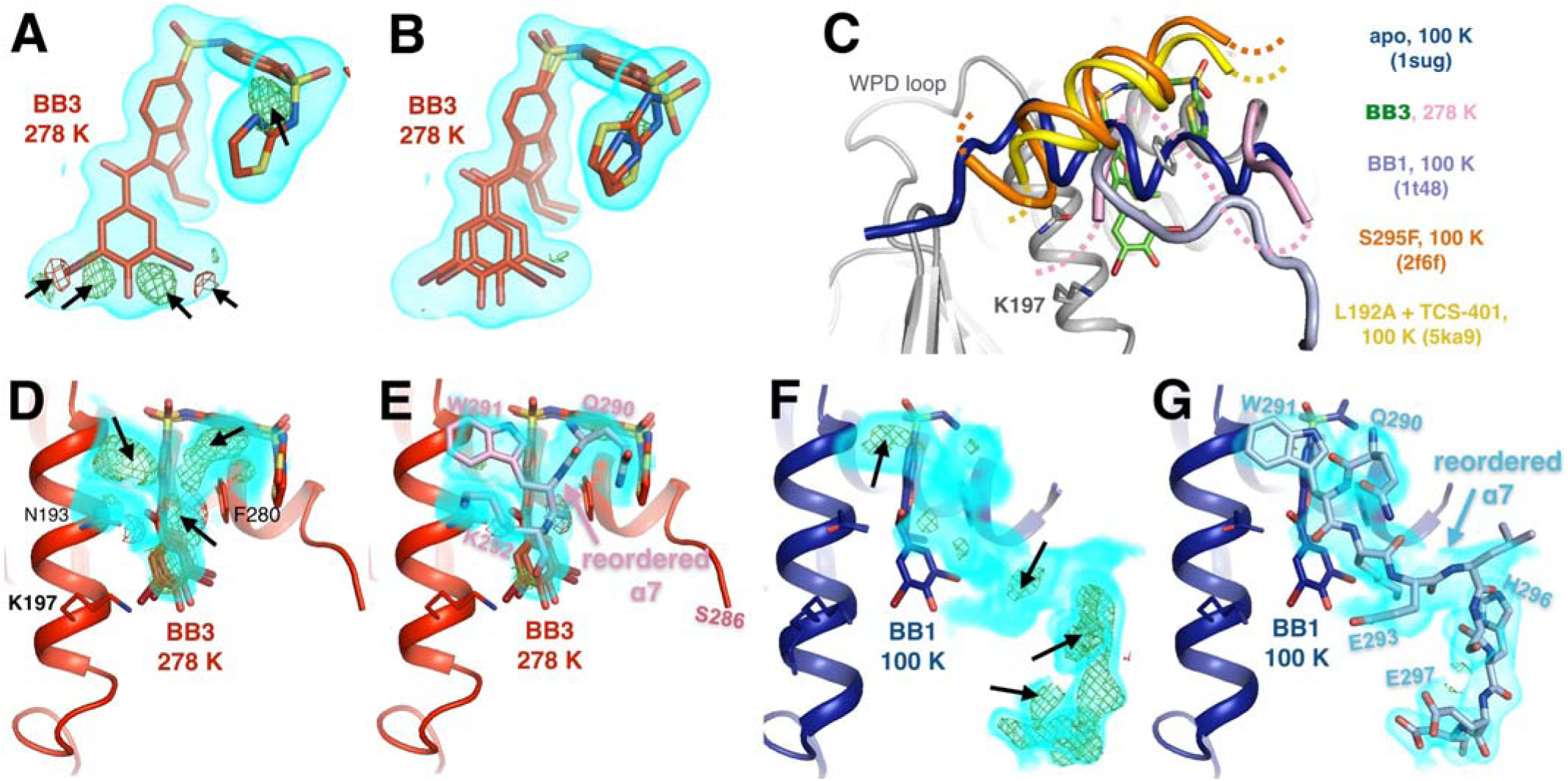
Allosteric-inhibitor-bound PTP1B has some low-occupancy conformations only at room temperature. **(A)** At room temperature, despite a good fit to 2Fo-Fc electron density, shown contoured at 0.75 σ (cyan volume), the allosteric inhibitor BB3 cannot be sufficiently modeled with a single conformer, as evidenced by +3.5 σ (green mesh) and -3.5 σ (red mesh) Fo-Fc difference electron density. These features are absent from the cryogenic structure with BB3 (1t4j). **(B)** These unexplained difference features disappear when a second alternate conformer is added with a translation at the “bottom” of the molecule (from this viewing angle) and dihedral-angle changes at the “top right”. **(C)** The disordered α7 helix is reordered above the BB binding site in various structures: room-temperature with BB3 (pink; BB3 molecule in green), cryogenic with BB1 (pale blue, PDB ID 1t48), cryogenic with the S295F mutation (orange, PDB ID 2f6f), and cryogenic with the L192A mutation and the TCS-401 active-site inhibitor (yellow, PDB ID 5ka9). The normal ordered α7 conformation (blue, PDB ID 1sug) is shown for reference. See **Figure 6A** for another example of a reordered α7 conformation. **(D-G)** Evidence for some of the conformations in **(C)**. **(D)** At room temperature, 2Fo-Fc electron density contoured at 1.0 σ (cyan volume) and Fo-Fc difference electron density contoured at +3.5 σ (green mesh) and -3.5 σ (red mesh) suggest something remains unmodeled above the bound BB3 molecule. **(E)** Modeling a reordered portion of the disordered α7 helix, including Trp291 (pink), fits the 2Fo-Fc density, removes the positive Fo-Fc peaks, and has reasonable interactions with nearby sidechains on α3 (left). **(F-G)** At cryogenic temperature, the electron density (shown at the same contour levels) suggests a different reordered conformation of α7 above BB1 (PDB ID 1t48). Both reordered α7 conformations (**D-E** vs. **F-G**) place residues 290-292, including Trp291, in the same place. However, the C-terminal portion of the reordered conformation with BB1 at 100 K, including His296 (right), is sterically incompatible with the N-terminal portion of the α6-α7 junction with BB3 at 278 K, including Ser286. Neither conformation is compatible with the electron density for the other, suggesting that differences in temperature and/or inhibitor dictate different α7 conformations.

### Multitemperature crystallography of loop 16 allosteric site

We also observed temperature dependent ordering in a loop (loop 16, L16; residues 237-243) that sits underneath the α6-α7 junction just beyond the BB binding site. At higher temperatures, the electron density (**Figure 3B**) clearly reveals that L16 adopts an alternative conformation with its backbone shifted by >5 □ from the primary conformation (**Figure 3D**). Modeling this alternative loop conformation back into the lower-temperature models and refining its occupancy reveals a temperature dependence (**Figure 3E, Figure 3 - Figure S1**) that is qualitatively similar to the temperature dependences of the WPD loop. Remarkably, this L16 alternative conformation sampled by apo PTP1B beautifully matches the L16 conformation when PTP1B is allosterically inhibited by BB2 (**Figure 3C**). This rearrangement provides further evidence that BB2 selects pre-existing, globally dispersed conformations rather than inducing new ones. This loop was not identified as part of the allosteric network in PTP1B based on NMR and MD studies [11]. However, it is seemingly coupled to the α6 helix: Lys239 from L16 H-bonds with Ile281 from α6 in the global closed state, but not the global open state in which L16 adopts its alternative conformation. Because α6 is directly coupled to the α7 order-disorder transition, we therefore propose that L16 is a newly recognized component of the collective allosteric network in PTP1B. Interestingly, an approach combining molecular dynamics and machine learning recently pointed to this area as a potential “cryptic” binding site [23]. Moreover, several residues lining the L16 site (Met3, Lys237, Ser242, Met282) were implicated as being allosterically linked to active site based on NMR chemical shift perturbations upon mutation of the WPD loop [24]. Therefore, the L16 site may be not only energetically coupled to the active site, but also capable of forming a new small-molecule binding pocket via the conformational heterogeneity we observe.

**Figure 3:**
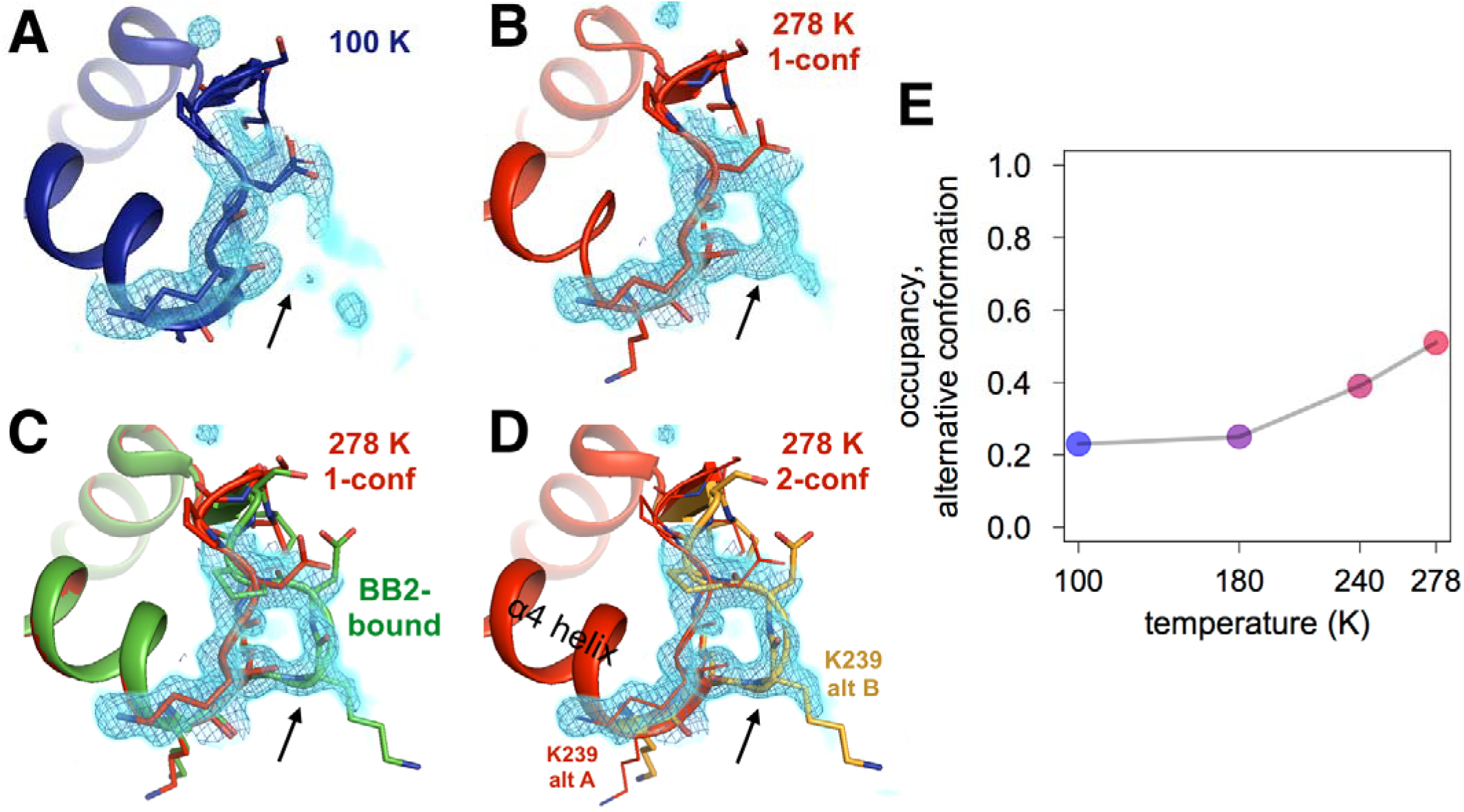
Both an allosteric inhibitor and high temperature favor an alternative conformation for an α7-coupled loop 16. **(A)** At low temperature, loop 16 (residues 237-243, bottom right) is single-conformer, as evidenced by 2Fo-Fc electron density contoured at 1.0 σ (cyan volume) and at 1.0 σ (blue mesh). **(B)** At high temperature, when the protein is modeled as single-conformer, the electron density suggests the existence of an alternative conformation. **(C)** The structure with BB2 bound (>12 □ away) (PDB ID 1t49) perfectly explains the mysterious electron density. **(D)** The final 278 K dual-conformation model is a good explanation of the data. **(E)** The refined occupancy of the alternative conformation (state “B”) in apo PTP1B increases continuously but non-linearly with temperature.

**Figure 3 - Figure S1:**
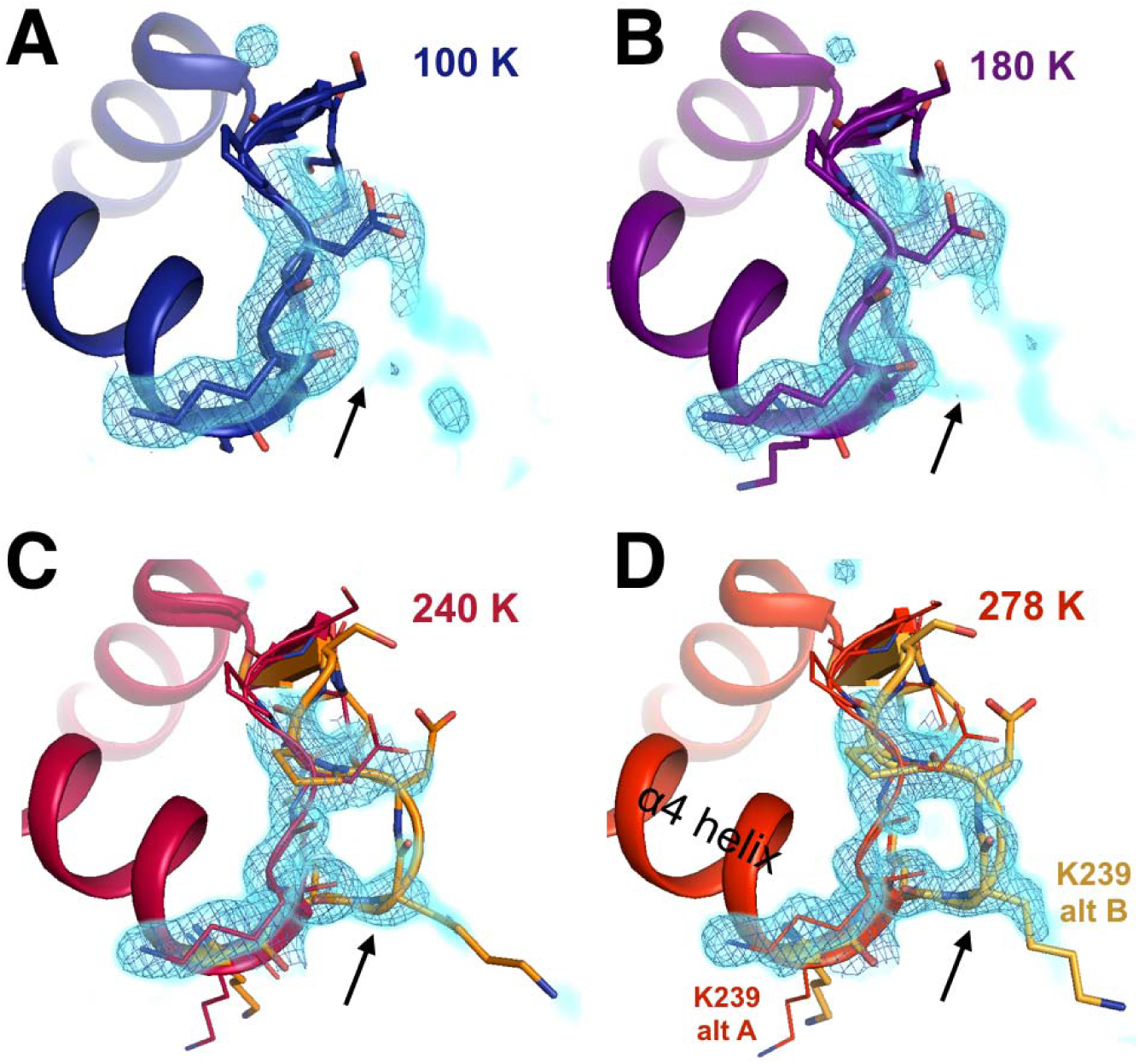
The conformational distribution of the α7-coupled loop 16 titrates with temperature. The loop consisting of residues 237-243 (bottom right) is single-conformer at low temperatures including **(A)** 100 K and **(B)** 180 K, as evidenced by 2Fo-Fc electron density contoured at 1.0 σ (cyan volume) and at 1.0 σ (blue mesh). By contrast, it adopts multiple conformations at higher temperatures including **(C)** 240 K and **(D)** 278 K.

### Multitemperature crystallography of 197 allosteric site

### Figure 4 - Movie S1

Movie version of **Figure 4B**.

In addition to the temperature-dependent conformational heterogeneity observed at and around the BB binding site, we observed residues with temperature-sensitive conformational heterogeneity in the “197 site” (**Figure 4**). Moreover, the alternative conformations of several residues in this region have a pattern of steric incompatibility with multiple states of the WPD loop and α7 helix, suggesting that the 197 site may be mechanistically linked to the active site in a similar way as the BB binding site.

**Figure 4:**
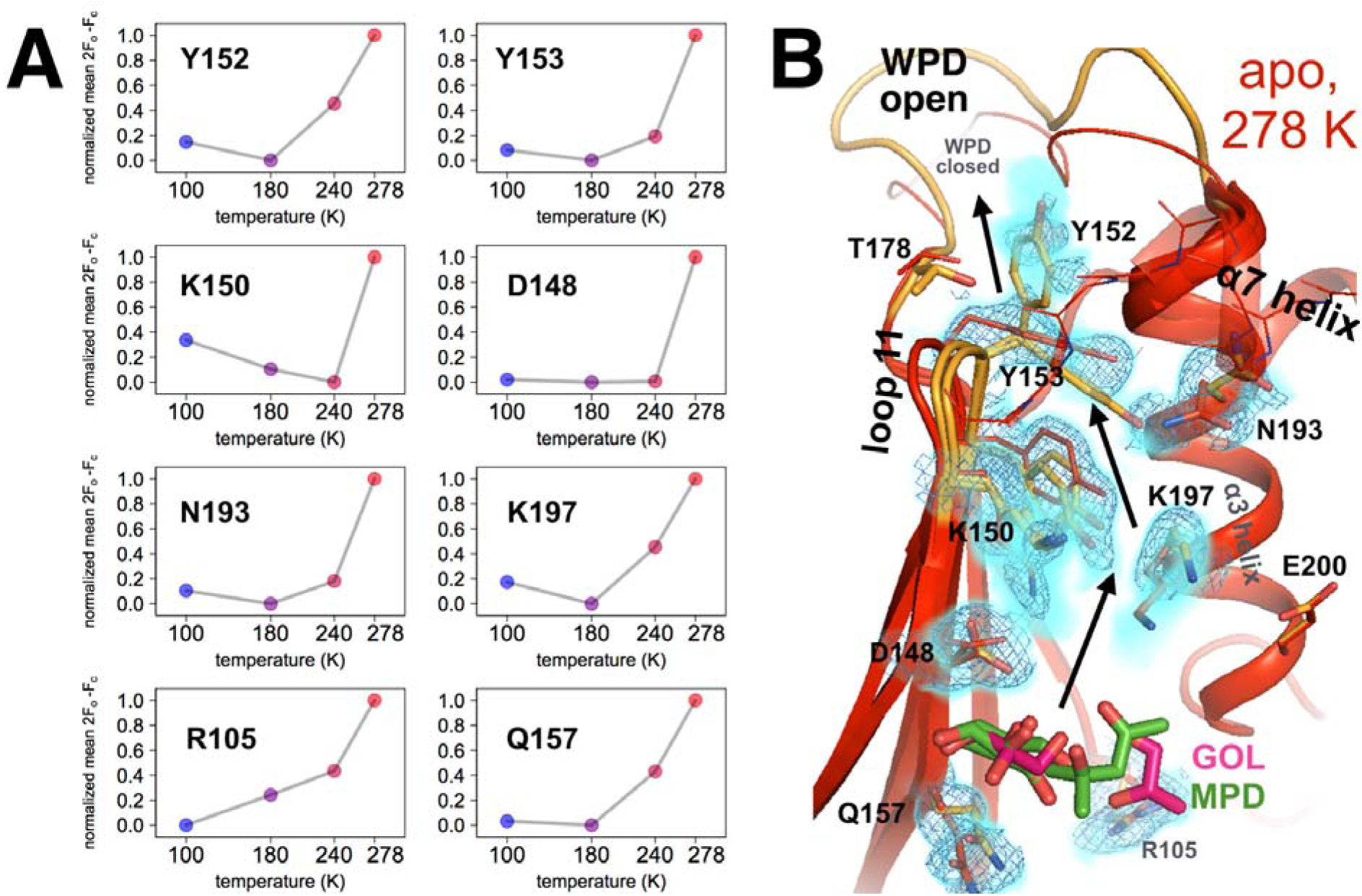
Coupled conformational heterogeneity leads to the new 197 allosteric site. **(A)** Several residues distinct from both the active site and previously characterized allosteric regions each have minor alternative conformations that become more populated with temperature. This is quantified by the sum of 2Fo-Fc electron density values for the centers of atoms that are unique to the minor state (defined as being at least 1.0 □ away from any atoms in the major state), normalized across temperatures from 0 to 1 for each residue. **(B)** These residues colocalize to a region of the protein surrounded by loop 11 (top-left), the quasi-ordered α7 helix (top-right), and the α3 helix (right), including the eponymous K197. 2Fo-Fc electron density contoured at 0.6 σ (cyan volume) and at 0.8 σ (blue mesh) justify multiple conformations for these residues in our 278 K apo model, as quantified in **(A)**. The alternative conformations of these residues appear to interact with one another and thus may be allosterically coupled. Ordered crystallization mother liquor or cryoprotectant molecules (glycerols in pink, from the PDB and our structures, or MPD molecules in green, from the PDB) can be present at the terminus of this allosteric pathway, suggesting it may be amenable to binding other small molecules.

A major link between the WPD loop and the 197 site is Tyr152. When the WPD loop is closed and the α7 helix is ordered, Tyr152 adopts a “down rotamer” (**Figure 4 - Figure S1**, red). By contrast, when the WPD loop is open and the α7 helix is disordered, the room-temperature electron density suggests that Tyr152 adopts an “up rotamer” (**Figure 4 - Figure S1C**, orange). However, difference electron density peaks remain (**Figure 4 - Figure S1C**) that indicate the presence of the down rotamer as an alternative conformation. Consistent with this interpretation, modeling just the additional down rotamer is insufficient to explain the density (**Figure 4 - Figure S1D**). These two rotamers are accommodated in the WPD-loop-open state by a shift of the L11 backbone (**Figure 4 - Figure S1D**). The down rotamer is sterically incompatible with phosphorylation of Tyr152, which occurs *in vivo* [25,26], suggesting that the up rotamer may have additional regulatory roles. Tyr152 in the L11 backbone conformation with just the down rotamer (red in **Figure 4 - Figure S1**) is sterically incompatible with the open WPD loop conformation (**Figure 4 - Figure S1E**). Similarly, the Tyr152 up rotamer is sterically incompatible with the ordered α7 conformation (**Figure 4 - Figure S1E**). In turn, α7 is conformationally synchronized with the WPD loop (**Figure 2A** and **Figure 1D**) and is a key hub connecting loop 11 and the WPD loop [11]. These results together suggest that the allosteric circuitry of PTP1B involving Tyr152 is complex and multibody. Tyr152 likely exemplifies a population shuffling mechanism whereby mixtures of microstates (rotameric state of Tyr152) exchange on a fast timescale as the protein transitions between macrostates (WPD loop state, α7 ordering, and L11 backbone shifting) on a slower timescale [27].

**Figure 4 - Figure S1:**
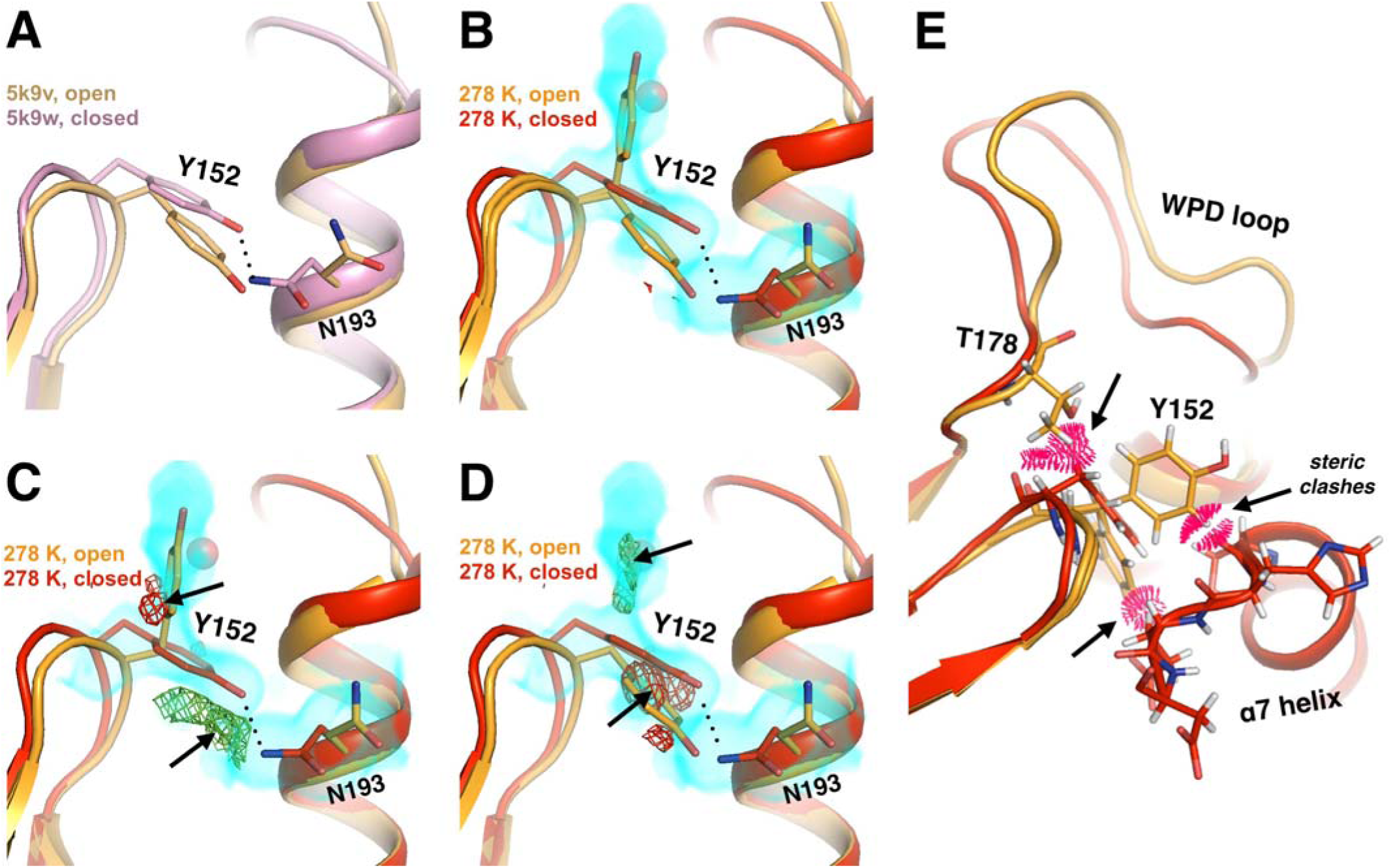
Alternative conformations in apo PTP1B recapitulate and expand upon reported coupling between loop 11 and α3. (A) Structures of the apo open state (yellow, 5k9v) and the closed state with the active-site inhibitor TCS401 (pink, 5k9w) [11] feature coupled movements of the loop 11 backbone, including Tyr152, and Asn193 within α7. **(B-D)** In our 278 K apo structure, we also observe coupling between these residues. However, Tyr152 is best fit with a “down” plus an “up” rotamer as alternative conformations for the open-state backbone, and only a down rotamer for the closed-state backbone. This is borne out by 2Fo-Fc electron density contoured at 0.3 σ (cyan volume) and Fo-Fc electron density contoured at +2.8 σ (green mesh) and -2.8 σ (red mesh) after refinement with either the down rotamer **(C)** or the up rotamer **(D)** omitted. A partial-occupancy water that is mutually exclusive with the up rotamer is also present (transparent red sphere). Note: Asn193 may also adopt another low-occupancy rotamer. **(E)** The down Tyr152 rotamer in the WPD-closed-compatible L11 backbone conformation (red) would sterically clash (pink pillows) with Thr178 in the open state of the WPD loop (orange). Both Tyr152 rotamers in the WPD-open state (orange) would sterically clash with the ordered conformation of the α7 helix (red).

In our non-cryogenic-temperature datasets, the electron density suggests a complex interplay between alternative conformations for Asn193 on the α3 helix and Tyr152 on loop 11 (L11) (**Figure 4 - Figure S1**). Chemical-shift-restrained molecular dynamics simulations also suggested that these residues have mutually coupled alternative conformations [11]. Asn193 is part of the α3 helix (residues 187-202), which immediately follows the WPD loop in sequence. The N-terminal region of this helix (through Phe196) rotates by 2-20°, resulting in shifts of 0.2-0.7 □ for some Cα atoms, based on cryogenic crystal structures of apo (WPD-open) vs. active-site-inhibitor-bound (WPD-closed) PTP1B [11]. Similarly, the multiconformer model for our 278 K apo dataset includes alternative backbone conformations for the WPD loop and the beginning of α3: through Asn193 plus Phe196-Lys197 (this is a conservative interpretation of which residues in the helix have alternative backbone conformations). Our results suggest that α3 inherently shifts as the protein transitions betweens its global macrostates even in the apo state.

Strikingly, several residues propagating down L11 from Tyr152, and down α3 from Asn193, also adopt multiple conformations at higher temperatures (**Figure 4**). These residues colocalize in a shallow pocket nestled between loop 11, the β4 and β5 strands, and the α3 and α7 helices. We will refer to this area here as the “197 site” because the sidechain of Lys197 extends into the pocket. Our analysis indicates a complex, inter-connected network involving multiple aromatic stacking, hydrogen-bonding, van der Waals, and electrostatic interactions. To complement this model-based assessment with a map-based approach, for several residues in the pocket we quantified electron density as a function of temperature for atom positions that are unique to the minor conformation (i.e., do not overlap with any atoms in the major conformation), reasoning that residues which respond to temperature similarly may be energetically coupled [12]. The population of each minor conformation increases non-linearly with temperature (**Figure 4A**) in a similar fashion as the open state of the WPD loop (**Figure 1D**) and the disordered state of the α7 helix (**Figure 2A**), in support of the idea that these various regions of the protein are mutually conformationally coupled. As with the L16 site, several residues at both ends of the 197 site (Tyr152, Tyr153, Lys150, Arg105) are suggested to be allosterically linked to the active site based on chemical shift perturbations [24].

We emphasize that the 197 site is structurally distinct from the previously characterized allosteric sites [7,10], so any small molecules that bind to this new site would represent a distinct strategy for inhibiting PTP1B. Surprisingly, in all of our multitemperature apo structures, ordered glycerols are present not only in the active site as mentioned above but also in the 197 site (**Figure 4 - Figure S2**), and MPD also binds here in another published structure (PDB ID 2cm2). These observations suggest that the 197 site may be bindable by other small molecules. We therefore hypothesized that binding of a small molecule to the 197 site could propagate changes in conformational heterogeneity to the WPD loop to interfere with catalysis.

**Figure 4 - Figure S2:**
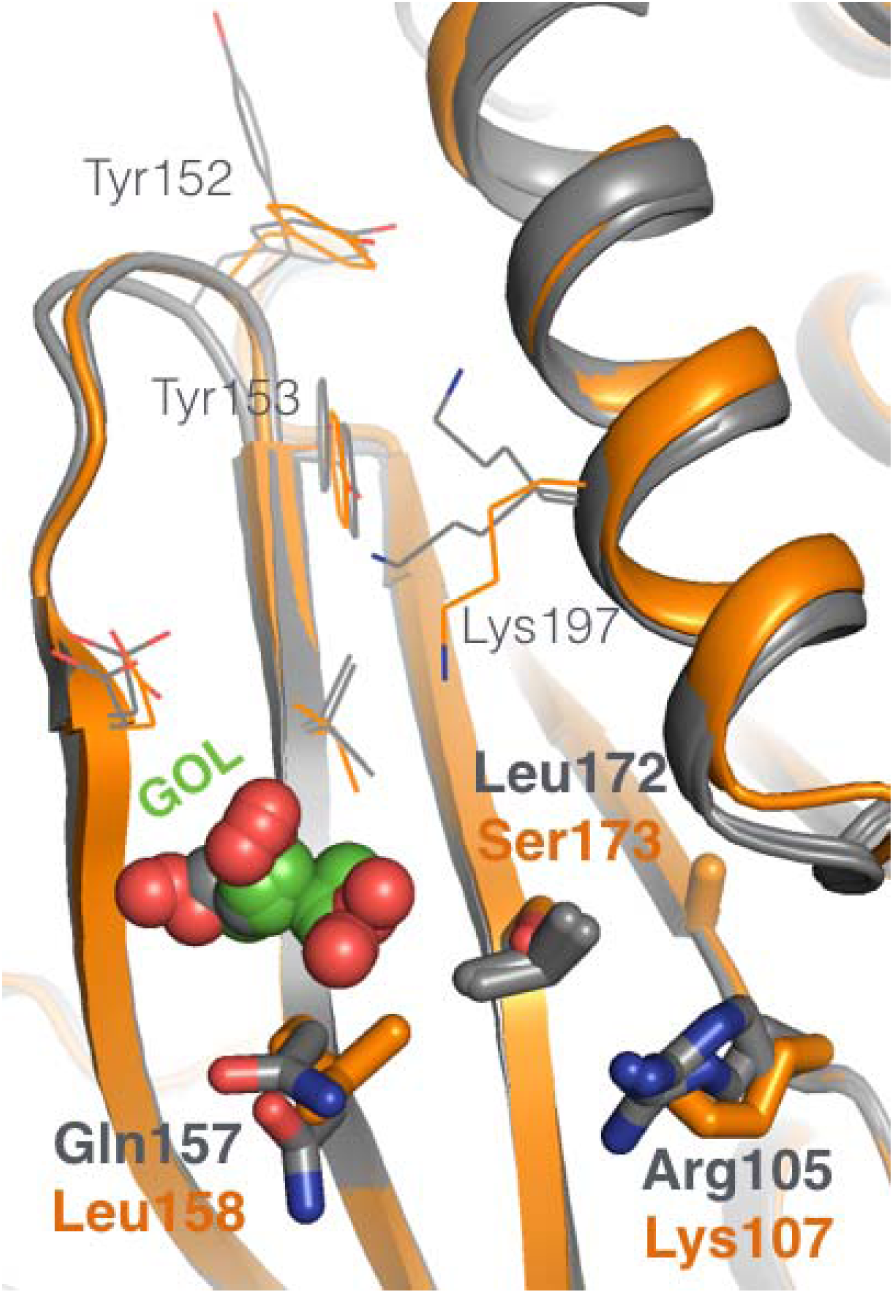
The new 197 allosteric site has local sequence differences in related PTPs. Several amino acids in the new 197 allosteric site in PTP1B (gray) are different at the equivalent positions in the closest homolog, TCPTP (orange).

To test whether more directed perturbations to this site can allosterically modulate enzyme function, we introduced “dynamically destructive” mutations (Y152G, Y153A, K197A) that were predicted to preserve the protein’s general structure, yet interfere with the conformational heterogeneity along the putative allosteric pathway lining the 197 site by removing interactions between alternative conformations. For Y152 we chose a mutation to glycine instead of alanine to more fully disengage residue 152 from the WPD loop, given that the Cβ and Hβ atoms of Y152 sterically engage with the WPD loop (**Figure 4 - Figure S1E**). All three mutations indeed reduce catalytic efficiency, to varying extents: the mutation nearest to the WPD loop (Y152G) reduced k_cat_/K_M_ the most, and the mutation farthest from the WPD loop (K197A) reduced k_cat_/K_M_ the least (**Figure 4 - Figure S3**). Our results are generally in line with reported effects for the Y152A+Y153A (“YAYA”) double mutation [11] and for the Y153A single mutation [24] despite differences in the precise protein construct being used. Overall, our results illustrate that local perturbations in the vicinity of the new 197 allosteric site can impact catalysis.

**Figure 4 - Figure S3:**
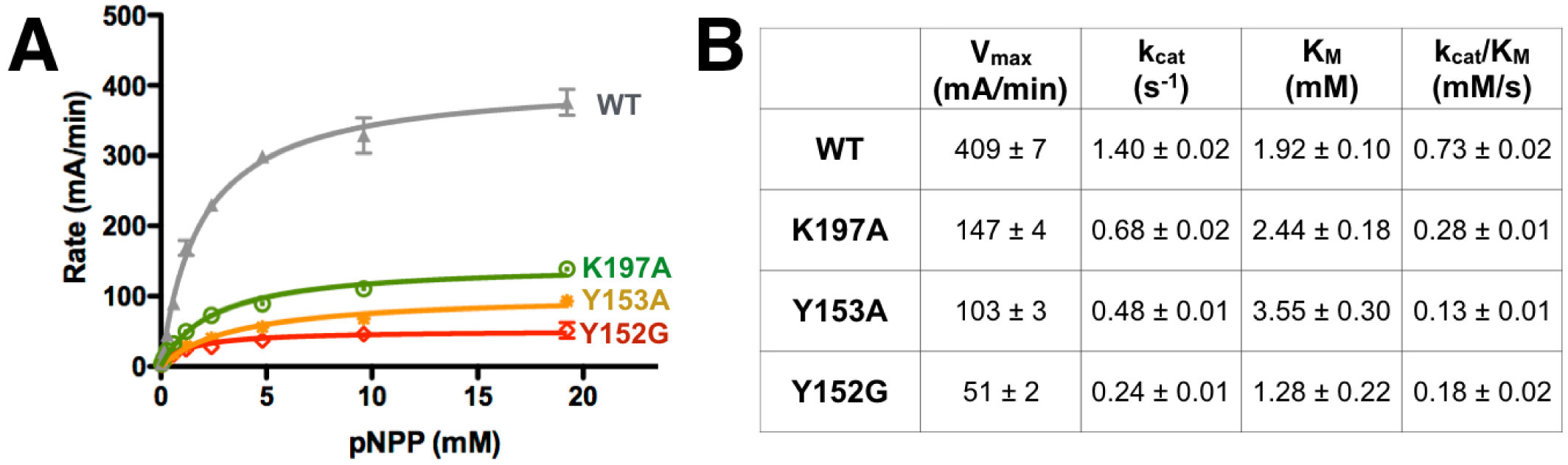
Mutations along the new allosteric pathway reduce enzyme activity. **(A)** Point mutations to several residues along the allosteric pathway from the new allosteric site to the active site reduce activity to varying degrees. Data represent the mean of four independent assays ± standard deviation. **(B)** Michaelis-Menten kinetics for WT vs. allosteric point mutants ± the standard error of the mean. See Methods for details.

**Figure 4 - Figure S4:**
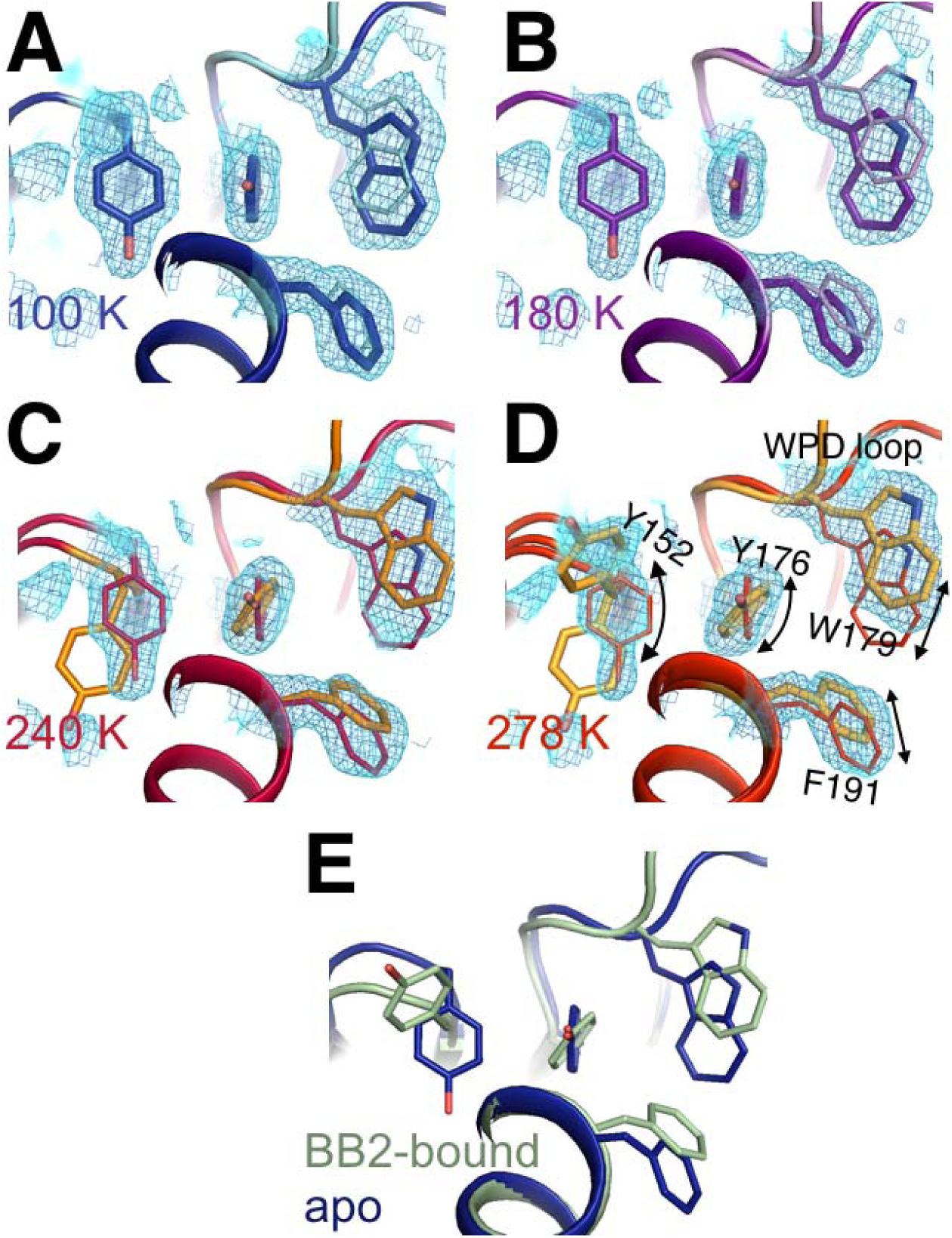
Flexible aromatic residues complete an allosteric circuit. Conformational heterogeneity for Tyr176 links Tyr152 in the new all`osteric pathway to Phe191 in the previously reported allosteric pathway [10] and Trp179 in the WPD loop, thus completing an allosteric circuit. **(A-D)** In apo PTP1B, 2Fo-Fc electron density contoured at 0.8 σ (cyan volume) and at 0.8 σ (blue mesh) generally justifies a single conformation these aromatic residues at low temperatures, but multiple conformations at high temperatures – especially for Tyr176 and Tyr152. **(E)** These individual conformations are evident in the previously published 100 K apo (blue, 1sug) or BB2-bound (green, 1t49) structures, respectively.

Overall, we describe a large, collectively coupled allosteric network on one contiguous face of the protein (**Figure 1E**). This network is interconnected not only on the surface, but also within the hydrophobic core. For example, Tyr176 adopts alternative sidechain conformations at higher temperatures that differ by a small rotation of the relatively non-rotameric χ2 dihedral angle [28] (**Figure 4 - Figure S4**). The two conformations of Tyr176 are structurally compatible with different conformations of the surface-exposed Tyr152 in one direction, and of the buried Trp179 in the WPD loop and BB allosteric pathway [10] in the other direction (**Figure 4 - Figure S4**). Thus, surface residues such as Tyr152 may be conformationally coupled to the buried underside of the WPD loop via a similar mechanism as BB binding – remotely modulating the Trp179 anchor via coordinated hydrophobic shifts – but from a different angle of attack, via Tyr176. Overall, such coordinated local shifts within the hydrophobic core likely “lubricate” the transition between discrete global states of PTP1B.

### Assessing the ligandability of the surface of PTP1B using automated crystallography

Although the results described above establish a conformationally coupled network within the structure of PTP1B, allosteric inhibition also requires binding sites for small molecules that can conformationally bias this network to modulate function. To identify potential allosteric ligand-binding sites in PTP1B, we mapped the small-molecule binding potential or “ligandability” of the entire protein surface. Specifically, we used small-molecule fragments, which by virtue of their small size provide a relatively large sampling of drug-like chemical space [29]. Astex Pharmaceuticals has previously explored fragment-based drug design on PTP1B [30]; however, that screen used molecules pre-selected to enrich for binders to phosphatase active sites, which contrasts with our goal of exploring the surface outside of the active site. To determine cocrystal structures of hundreds of fragments with PTP1B, we used the high-throughput fragment-soaking and crystallographic pipeline available at Diamond Light Source [13] to individually soak 1,918 apo PTP1B crystals with small-molecule fragments in DMSO from several curated libraries, and another 48 with just DMSO. We then used robotic sample handling to automatically collect complete X-ray datasets at cryogenic temperature (**Figure 5 - Table S1**). Of the 1,966 total soaks, 1,774 yielded diffraction data that could be successfully processed. The data were generally high-resolution: the average resolution was 2.1 □, 65% of resolutions were better than 2.0 □, and 87% were better than 2.5 □ (**Figure 5A, Figure 5 - Table S1**). The large number of datasets enabled us to use the new Pan-Dataset Density Analysis (PanDDA) algorithm [14] to reveal bound fragments. PanDDA performs weighted subtractions of the “background” electron density (computed from apo and unbound datasets) from each electron density map (**Figure 5B-C**). The optimal subtraction yields electron density corresponding to the ligand-bound fraction of unit cells in the crystal.

**Figure 5:**
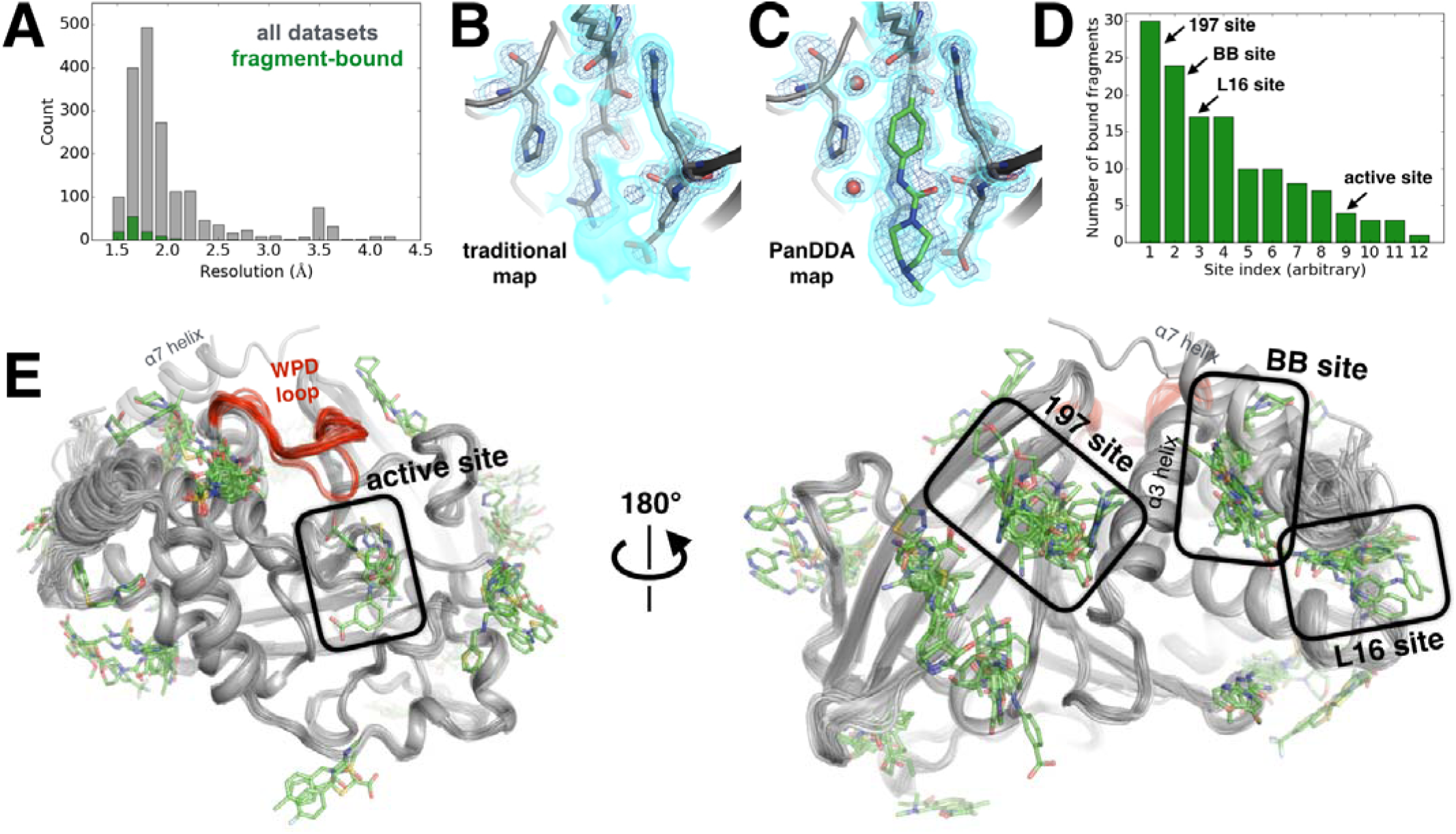
Electron-density background subtraction reveals small-molecule fragments at allosteric sites in PTP1B. **(A)** Histogram of X-ray resolution for 1,774 structures of PTP1B soaked with small-molecule fragments (gray) vs. the 110 structures from that set with small-molecule fragments bound to PTP1B (green). **(B)** For one example fragment, a traditional 2Fo-Fc map contoured at 1.25 σ (cyan volume) and at 3.5 σ (blue mesh) provides no clear evidence for a bound fragment. **(C)** By contrast, a background-subtracted PanDDA event map (85% background subtraction in this case) contoured at the same levels very clearly reveals the precise pose of the bound fragment, plus additional ordered water molecules that accompany it (red spheres). **(D)** PanDDA analysis and manual inspection reveal 110 fragment-bound structures of PTP1B, with bound fragments clustered into 12 non-overlapping binding sites. Some structures contain multiple bound copies of the same fragment. Several sites of interest are labeled. **(E)** Overview of bound fragments across the PTP1B surface. *Left*: front of protein, facing active site (WPD loop open and closed conformations in red). *Right*: back of protein, facing several fragment-binding hotspots: the 197 site, BB site, and L16 site.

#### Figure 5 - Table S1: Results of all 1,966 fragment and DMSO soaks into PTP1B crystals.

For each crystal soak of DMSO (only 48 soaks) or a fragment (1,918 soaks), we list temperature of data collection (almost always cryogenic), resolution of data set collected (with “None” for failed soaks), fragment name, SMILES string, fragment library, estimated occupancy of primary bound ligand from PanDDA (or “None” if no modelled ligand), and PDB ID. For a relatively small number of soaks, 2 fragments were soaked per crystal well (in 43 cases) or a database error led to uncertainty as to which of up to 16 fragments were soaked into the crystal (in 39 cases); in these cases, all possible fragments are listed. In 3 of these 82 cases we identified and modeled the ligand from among the possibilities based on the electron density in the event map. (available as: Figure 5 - Table S1.csv)

### Figure 5 - Movie S1

Movie version of **Figure 5E**.

Our PanDDA analysis of 1,774 datasets revealed 381 putative binding events. We manually inspected each putative binding event, and were able to confidently model the fragment in atomic detail for 110 hits (**Figure 5D**). Overall, 12 different sites in PTP1B were observed to bind fragments (**Figure 5E**). These sites are structurally distinct from one another – i.e., they share no residues in common, and fragments bound within different sites do not overlap with each other. They are also widely distributed across the protein surface. 25 fragments bind to multiple sites, but promiscuous binding is not unexpected from such small fragments, and still provides valuable information about favorable binding poses in each site.

PanDDA initially identified >80 putative events in the active site. Many of these can be attributed to movements of the WPD loop (**Figure 1**), often induced by oxidation of the catalytic Cys215, which is a natural regulatory mechanism [31]. Apart from these protein events and other false positives, we observe 4 fragments bound in the active site. This number is relatively low likely because our libraries were not customized to bind to the highly charged active site of PTP1B, as was the case in other studies [30].

To identify allosteric binders, we examined sites outside of the active site. Strikingly, we observed 24 bound fragments in the BB allosteric site (**Figure 6A**). The poses of many of these fragments overlap portions of the BB scaffold (**Figure 6A**). This result validates the idea that sites revealed by fragment screening may be fruitfully targeted for allosteric inhibition. In one structure with a fragment bound in this site, the α7 helix adopts a reordered conformation that covers the binding site (**Figure 6A**), reminiscent of other examples in published structures and in our high-temperature datasets (**Figure 2 - Figure S3**).

**Figure 6:**
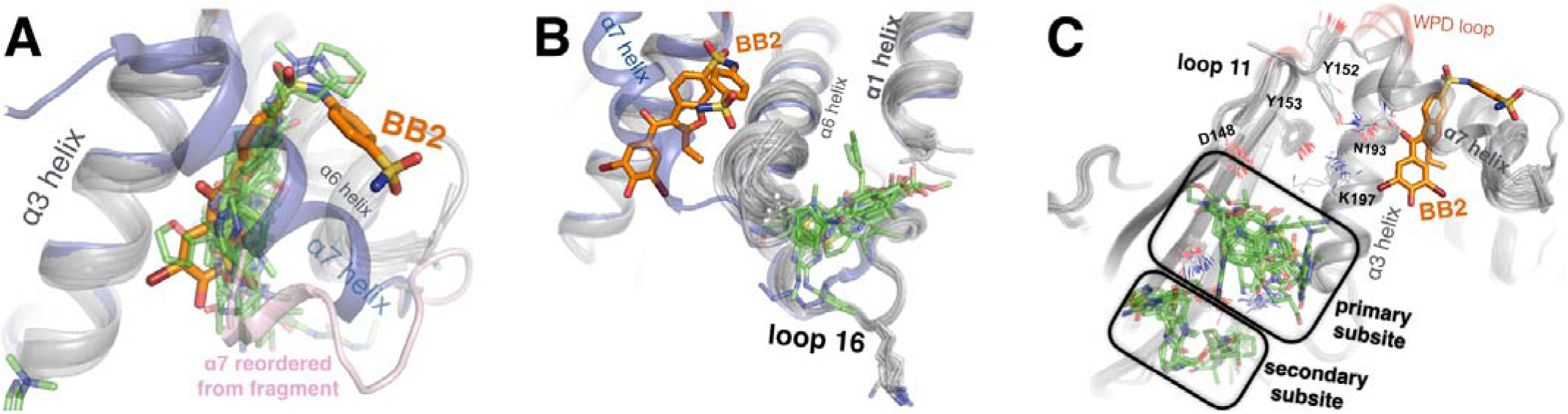
Fragments cluster at three binding hotspots distal from the active site. **(A)** 24 fragments (green) bind to the same site and in similar poses as the BB2 inhibitor (orange, PDB ID 1t49), and similarly displace the α7 helix (foreground, transparent blue, PDB ID 1sug). BB2 is also shown in the following panels to emphasize that its binding site is distinct from the other fragment-binding hotspots. One structure with a fragment bound in this site features a reordered conformation of the α7 helix (pink). **(B)** 17 fragments bind to the L16 site, where they may modulate the conformations of loop 16, the α6 helix, and the protein’s N-terminus on the α1 helix. **(C)** 30 fragments bind to the 197 site in one primary subsite contacting K197, or a distinct secondary subsite nearby.

**Figure 6 - Figure S1:**
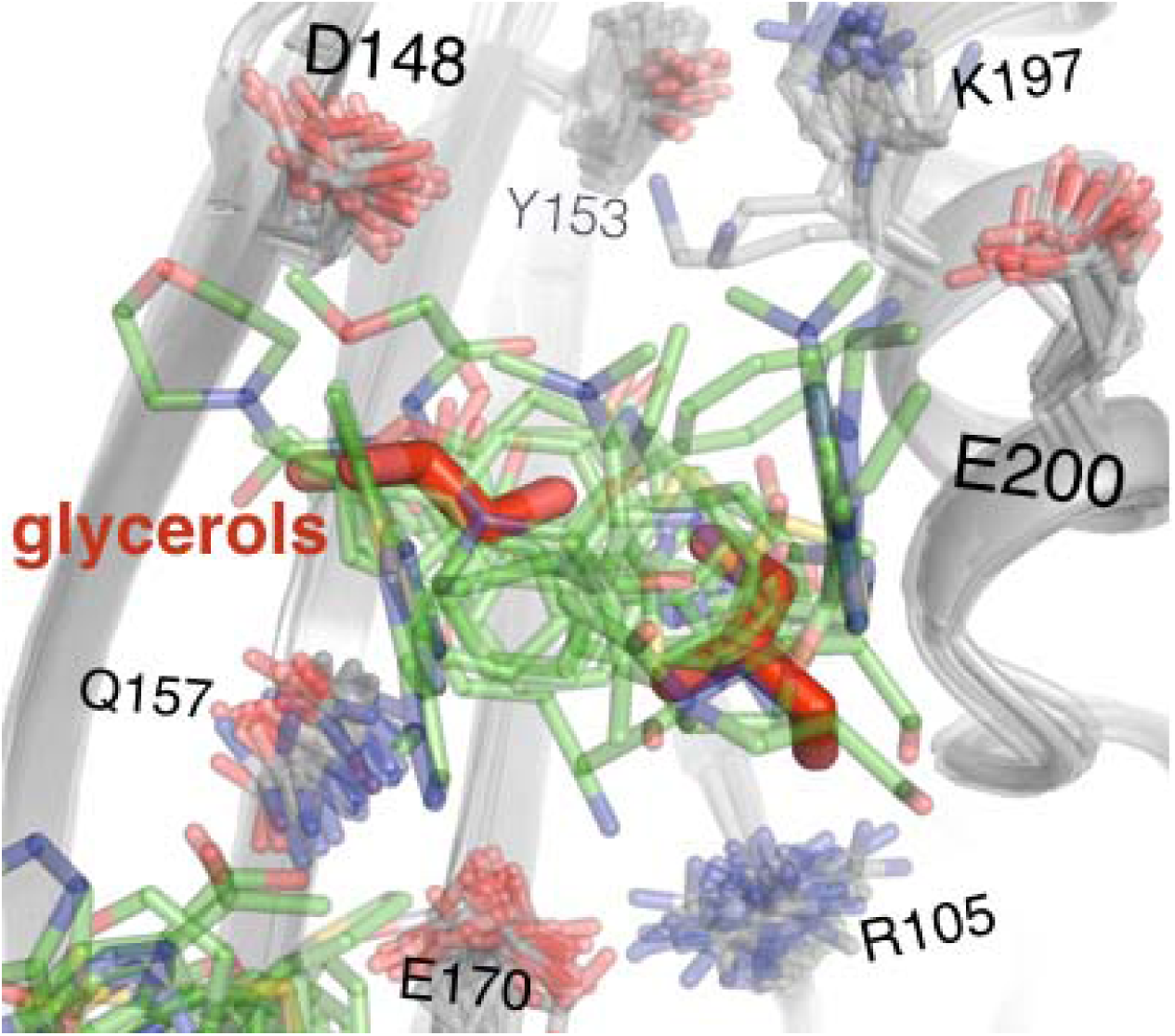
Fragments in the new 197 site overlay with glycerols from multitemperature structures. 15 fragments (green) in the primary subsite of the 197 site occupy a similar region of space as ordered glycerols (red) from our 278 K apo structure.

We also examined fragments bound to the L16 site and the 197 site, which were revealed as allosteric sites by our multitemperature analysis of apo PTP1B. Excitingly, both sites are fragment-binding hotspots: 17 fragments bind to the L16 site (**Figure 6B**) and 30 fragments bind to the 197 site (**Figure 6C**). Thus, distinct methods to gauge allosteric coupling and ligandability converge on the same regions of PTP1B.

The L16 site is between loop 16 (L16), the beginning of α1, and the end of α6. Most of the 17 fragments that bind here appear to “pry apart” these elements (**Figure 6B**) to create a cryptic binding site [23]. Because the end of α6 is coupled to the beginning of α7, which is perhaps the central allosteric hub of PTP1B [11], this site seems promising for allosteric inhibition. The fragments that bind here are diverse, but have some common features: aromatic moieties sandwich between Pro239 (of L16) and Met282 (α6), and carboxyl groups hydrogen-bond to the backbone amide of Glu2 (α6). These fragments do not spatially overlap with any fragments in the nearby BB site, confirming that the L16 site is genuinely distinct from the previously characterized allosteric site.

The 197 site is on the opposite side of the BB site, near α3 (including Lys197) and L11 (including Tyr152). 30 fragments bind in the 197 site, with 14 in the primary subsite near Lys197, and 17 in a nearby but distinct subsite separated by a “ridge” formed by the Gln157 and Glu170 sidechains (**Figure 6C**) (one fragment binds in both the primary subsite and the secondary subsite). These fragments are characterized by packing of aromatic moieties above Leu172, with additional aromatic or polar extensions in various directions. As with the L16 site, fragments in this site do not overlap with any fragments in the nearby BB site. However, several of the fragments in the 197 site do overlap with the positions of ordered glycerols (which were absent from all fragment-soaked structures to avoid competition for binding) from our multitemperature structures (**Figure 6 - Figure S1**). Similarly, glycerol in PDB ID 3qkp and β-octylglucoside in PDB ID 2cmc (among other examples) bind to sites that are occupied by fragments in our structures. These findings emphasize that fortuitous binding of buffer components and other miscellaneous compounds can in some cases provide useful information about binding sites [32]. It may be possible to link fragments in the primary subsite and secondary subsite to increase binding affinity. Although some fragments in the secondary subsite are largely stabilized by crystal-lattice contacts, they still enjoy favorable interactions with the protein that could potentially be useful for fragment extension. By contrast, the primary subsite is generally free from crystal-lattice contacts.

To assess the effect of the bound fragments on the structure of PTP1B more globally, for each dataset we built an ensemble structure consisting of both the ground state and the bound state. Each dataset was modeled with an innovative PDB format as a multiconformer structure that represents both a heterogeneous apo state and a heterogeneous holo state. Due to limitations in the PDB model format and the ability of conventional refinement programs to interpret and create reasonable restraints for this model type, either one conformation or four alternative conformations were used to describe each residue, often when only two were necessary. Due to this forced degeneracy, refinement of coordinates, occupancy, and B-factors must be highly restrained. We interpret the resulting occupancies as a good approximation of the fraction of unit cells that have a ligand present. Refining these ensemble structures using restraints that avoid overfitting allowed for some structural differences between the two state to emerge. In principle, these structural differences could give some prediction of the functional effects one might expect upon developing a higher-affinity version of the molecule. The refined ensemble structures were of high quality (**Figure 6 - Table S1**). However, generally speaking, the structural differences were subtle: the global backbone RMSD (N, Cα, C atoms) between the ground state and bound state ranged from 0.7-1.7 □. Cases with larger RMSD (>1.25 □) generally involved either active-site fragments that directly shift the WPD loop, or fortuitous oxidation of the active-site Cys215 [31]. Thus, fragment binding did not dramatically shift PTP1B from the open to the closed state in many these structures. Many of these fragments are certainly benign binders that bind to non-allosteric sites. However, the strong preference for the open state even with fragments that bind to allosteric sites is likely due to the absence of glycerol, which is present in our multitemperature structures (see Methods). Weak fragments do not overcome this energetic preference, and instead elicit conformational changes primarily in their immediate vicinity. Including glycerol to place the protein in a regime in which the open vs. closed states are more nearly isoenergetic during fragment soaks could potentially interfere with fragment binding to the 197 site, since ordered glycerols also fortuitously bind there (**Figure 6 - Figure S1**).

#### Figure 6 - Table S1: Crystallographic statistics for fragment-bound structures.

Resolution, R_work_, R_free_, and MolProbity score are listed for the 110 fragment-bound ensemble structures. (available as: Figure 6 - Table S1.csv)

### Increased occupancy via covalent tethering to test for allosteric inhibition

To test for allosteric effects from fragments in the 197 site or L16 site, we performed enzyme activity assays with 20 fragments that were thought to bind in either site during early rounds of iterative PanDDA analysis (see Methods) (Figure 6 - Table S2). We saw no inhibition of enzyme activity by the fragments, even at the maximum concentrations we were able to assay due to solubility of the ligands. This is not surprising given the fragments’ low affinities: soaking with fragments at 30-150 mM concentrations resulted in observed occupancies of only 10-30% in crystals. Medicinal chemistry efforts to improve the affinities of molecules that bind in the 197 and L16 sites by fragment linking and other elaborations represent a promising future direction. However, here we focus on an alternative strategy to validate the concept of allosteric inhibition at the 197 site: covalent tethering to enhance ligand occupancy.

**Figure 6 - Table S2:**
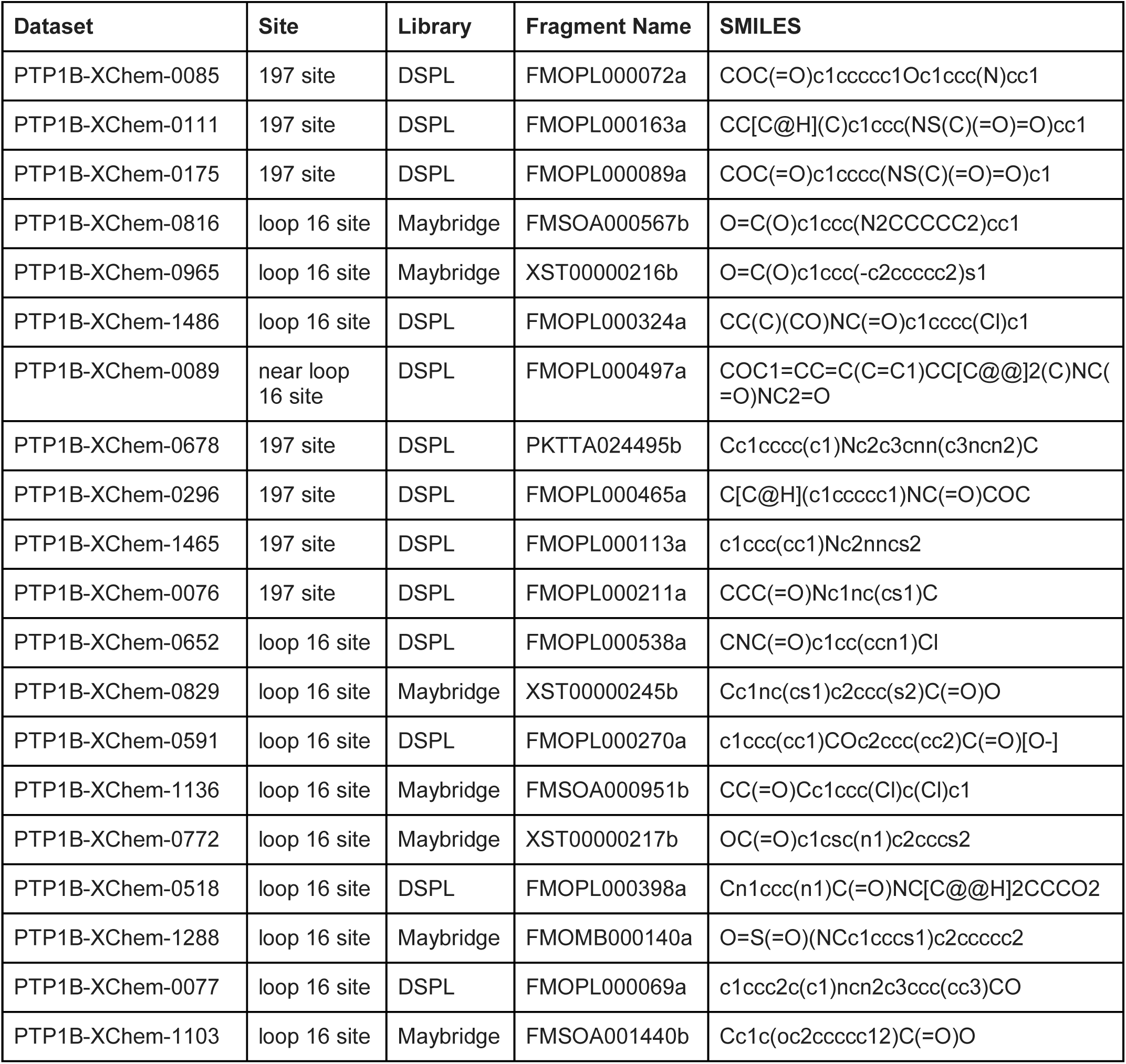
Small-molecule fragments tested in enzyme inhibition assays. These 20 small-molecule fragments, which were thought to bind in or near either the 197 site of the loop 16 site in our structures early in our PanDDA analysis, were tested for inhibition of PTP1B using a pNPP activity assay in quadruplicate with 200 nM protein and 1 mM fragment in 2% DMSO (final). No inhibition was observed in any case, compared to no-fragment and no-protein controls.

Specifically, we used “Tethering” [15,33] in which a residue near the site of interest is mutated to cysteine, then the mutant is mixed with disulfide fragments under partially reducing conditions. Affinity of the fragments for the site of interest drives the formation of a disulfide bond between the fragment and the adjacent cysteine. The extent of cysteine labeling can be measured using whole-protein mass spectrometry, and serves as a metric to rank the affinity of fragments for a given site. One major advantage of Tethering over other fragment-based approaches is that it can leverage low-affinity binding events into quantitatively labeled protein species, whose enzymatic activity can then be assayed.

For the new 197 allosteric site in PTP1B, we chose to tether to a K197C mutant for several reasons. First, K197 is on the α3 helix, which is a key allosteric element in PTP1B [11]. We predicted that small-molecule tethering to our site could could perturb the helix via K197C, perhaps mimicking the effects of a free molecule binding to the WT protein and altering the K197 conformational distribution. Second, K197 and E200 are the two residues on α3 whose Cα-Cβ vectors point in roughly the correct direction toward our new allosteric site. However, E200 engages in crystal-lattice contacts which would interfere with tethering in our P3_1_21 space group, so we focused on K197C instead.

To efficiently explore the chemical space of covalent small molecules for the 197 site, we used a library of 1,600 disulfide-capped fragments designed for covalent tethering experiments [34,35]. From our initial screen we identified 50 fragments that tethered to K197C >3 standard deviations above the average percent tethering for all 1,600 compounds (**Figure 7 - Figure S1A**). We next measured the ability of these top fragments to modulate PTP1B’s phosphatase activity (**Figure 7 - Figure S1B**). Formation of the tethered complex followed by a pNPP assay identified only one fragment, **1** (**Figure 7 - Figure S1C**), that appeared to inhibit PTP1B at a percentage comparable to the percentage of tethered complex (**Figure 7 - Figure S1B**), suggesting a direct relationship between labeling and inhibition. While **1** showed the activity we desired, the percent labeling and inhibition were relatively low. We hypothesized that altering the linker between the fragment core and the disulfide bond may lead to improved interactions between the protein and small molecule. For this reason we designed and synthesized **2** (**Figure 7A**), which has the orientation of the amide bond reversed, allowing for one less carbon in the disulfide linker (**Figure 7 - Figure S1C**).

**Figure 7:**
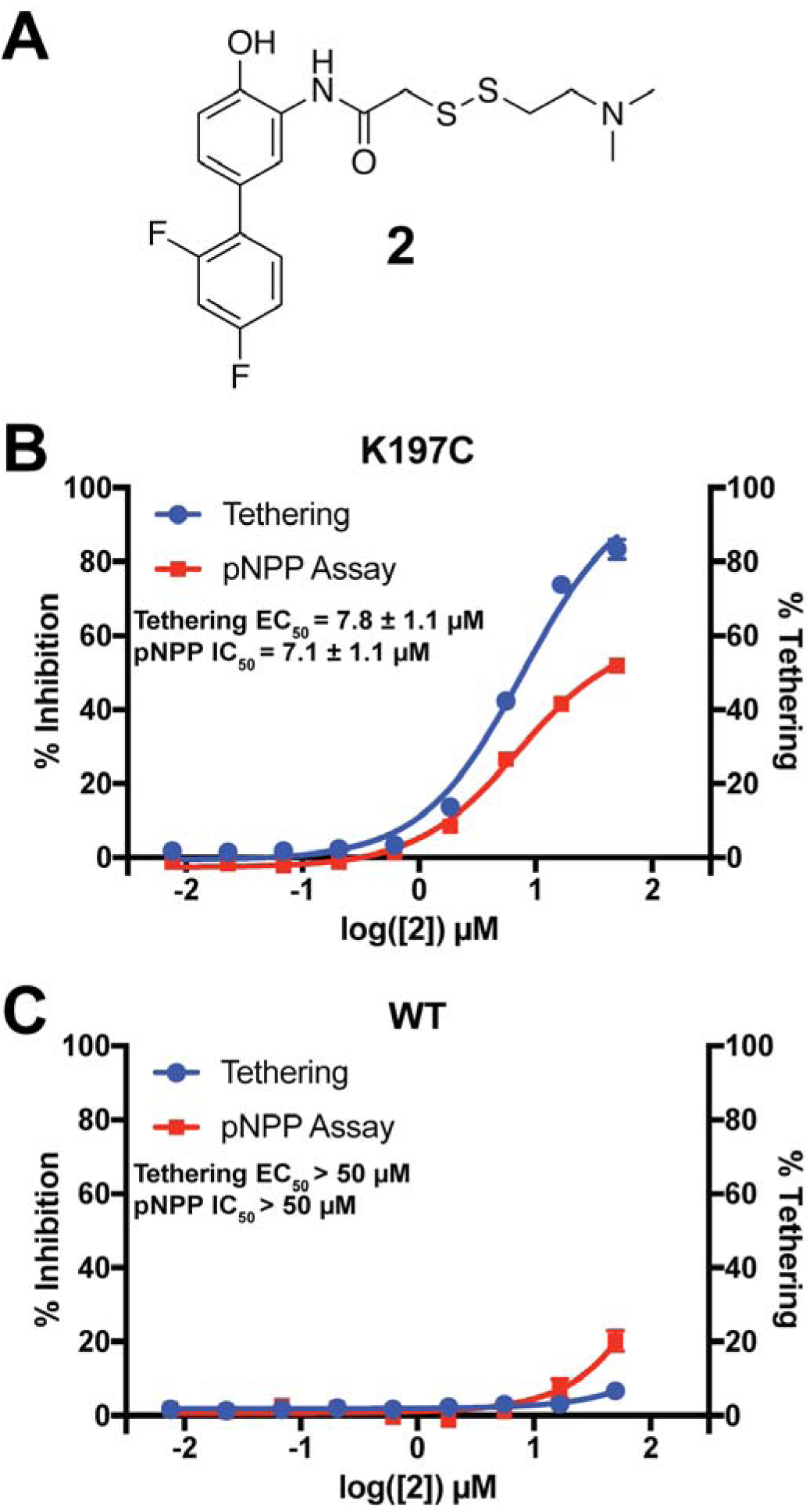
Characterization of a functional covalent allosteric inhibitor. **(A)** The chemical structure of our covalent disulfide fragment **2**. **(B)** Tethering and Inhibition of K197C at varying concentrations of **2**. The tethering EC_50_ observed was 7.8 ± 1.1 µM and the IC_50_ for pNPP activity was 7.1 ± 1.1 µM with a maximum inhibition of ~60%. Tethering data represents all tethering events combined. **(C)** Tethering and Inhibition of WT at varying concentrations of **2**. The tethering EC_50_ and the IC_50_ for pNPP activity were both > 50 µM. Tethering data represents all tethering events combined. Data represent the mean of three independent assays ± the standard error of the mean. All assays were performed in the presence of 100 µM of β-mercaptoethanol.

When assayed, **2** showed improved tethering and inhibition of K197C relative to **1**. **2** exhibited dose-dependent tethering and inhibition of K197C with a tethering EC_50_ of 7.8 ± 1.1 µM and an IC_50_ for pNPP activity of 7.1 ± 1.1 µM (maximum inhibition of ~60 µM) (**Figure 7B and Figure 7 - Figure S2A**). Importantly, **2** appeared to show little to no tethering of WT and minimal inhibition, supporting that **2**’s activity is specific to the K197 site and not due to tethering of the active-site cysteine found in both K197C and WT (**Figure 7C and Figure 7 - Figure S2B**). In fact, the inhibition that is observed for WT does not correlate with tethering, suggesting the inhibition may be from nonspecific factors, such as aggregation, at higher concentrations of **2** (≥ 50 µM). Michaelis–Menten kinetic analysis of K197C in the presence **2** (50 µM) showed a ~50% reduction in V_max_ relative to DMSO treatment, with little effect on the K_M_ for pNPP substrate (**Figure 7 - Figure S2C**). This supports a non-competitive allosteric mechanism of inhibition. The effect on WT kinetics was similar to the nonspecific inhibition observed in the dose titration experiment (**Figure 7 - Figure S2C**), once again supporting that the activity of **2** is specific for the K197 site on PTP1B.

**Figure 7 - Figure S1:**
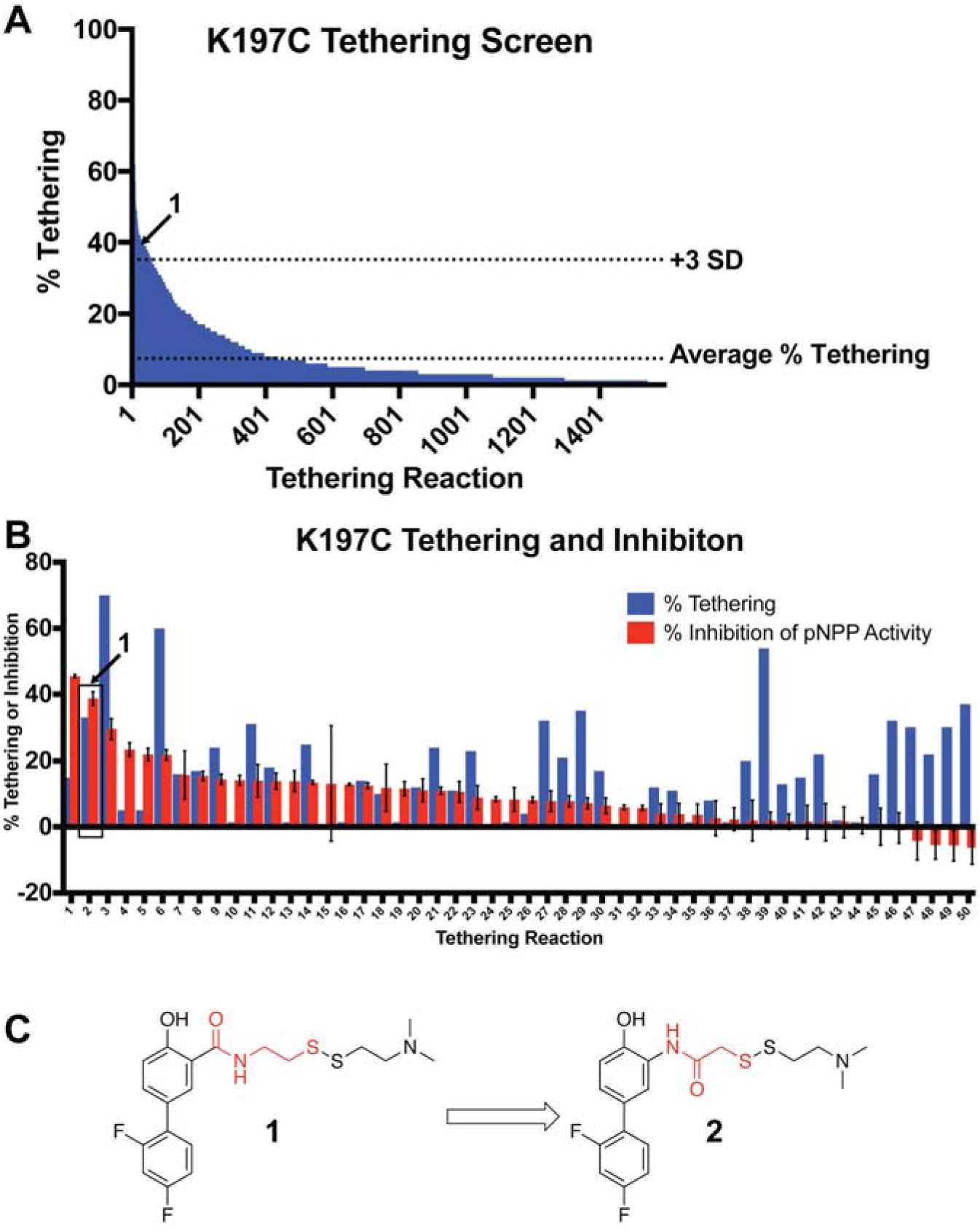
Identification of a functional covalent allosteric inhibitor. **(A)** 1,600 disulfide fragments were screened for their ability to covalently label K197C. The location of **1** in the data is highlighted. Data represents a single experiment. The % Tethering data reported is for the 1:1 (**2**:K197C) adduct only. **(B)** Follow up tethering and pNPP inhibition assays of the top 50 fragments from the initial screen. **1**, showed the second best inhibition of pNPP activity and correlated well with percent tethering. This suggested the activity was specific through K197C. Other fragment tested showed either poor, or non-specific inhibition of pNPP activity. Percent tethering represents a single experiment. The % Tethering data reported is for the 1:1 (**2**:K197C) adduct only. Percent inhibition represents the average of three independent assays ± the standard error of the mean. Both assays were performed in the presence of 50 µM of **1** and 100 µM of β-mercaptoethanol. **(D)** Structure of **1** as well as the closely related analog **2**. Molecular differences between **1** and **2** are highlighted in red.

**Figure 7 - Figure S2:**
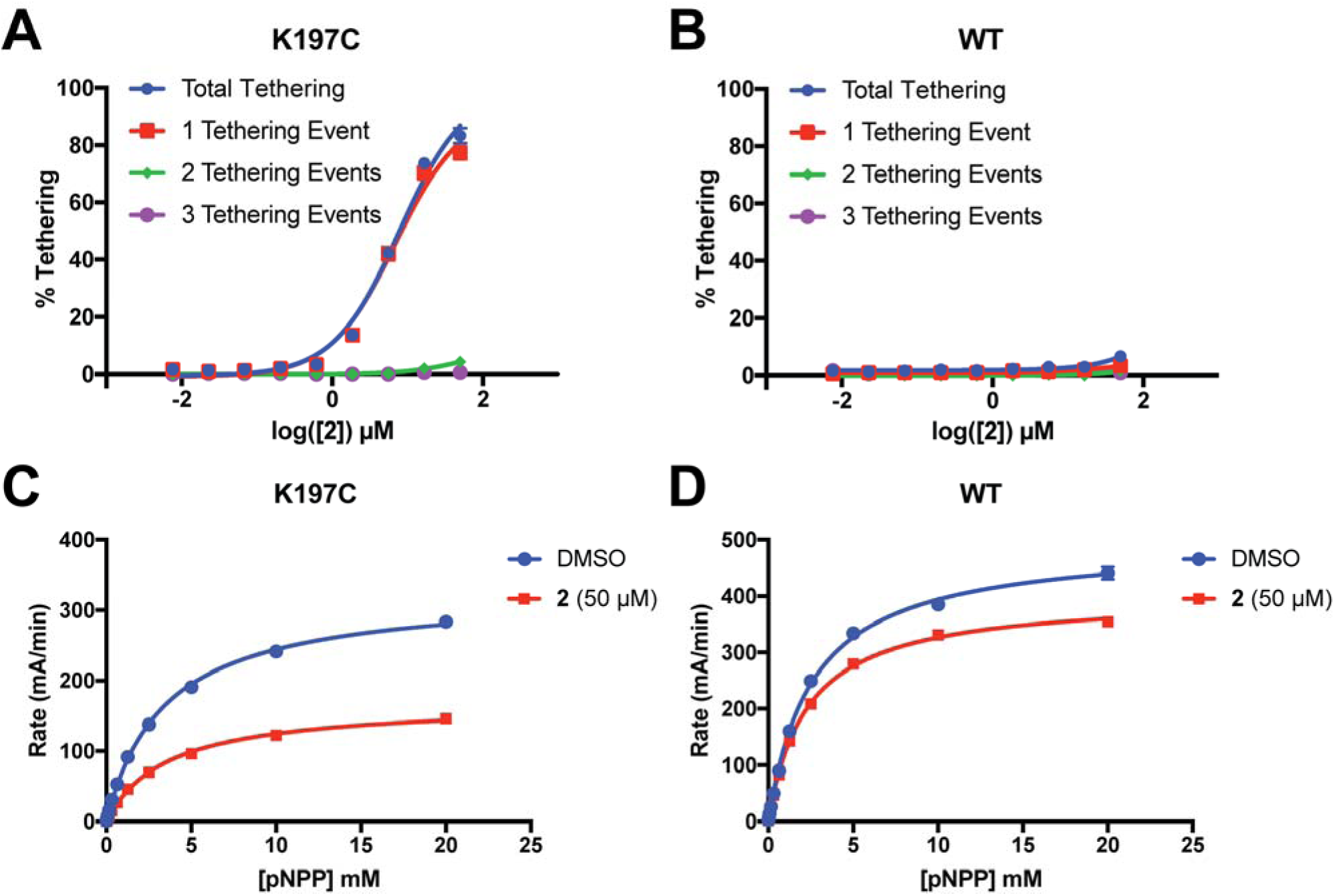
2 tethers to only a single cysteine in K197C and inhibits through a reduction in V_max_. **(A)** Percent tethering of K197C at varying concentrations of **2**. The vast majority of labeled K197C appears to have only a single tethering event of **2**, supporting specific labeling of K197C. Observation of more than one tethering event suggests nonspecific labeling of native cysteines in PTP1B. **(B)** Percent tethering of WT at varying concentrations of **2**. Very little tethering of WT is observed (<10%) with the total coming from a combination of different adducts, supporting nonspecific labeling. **(C)** Michaelis–Menten kinetic analysis of K197C in the presence and absence of **2** (50 µM). Compound **2** causes a ~54% reduction in V_max_ relative to DMSO but almost no change in K_M_, supporting a non-competitive allosteric mechanism of inhibition. **(D)** Michaelis–Menten kinetic analysis of WT in the presence and absence of **2** (50 µM). Compound **2** causes only a ~20% reduction in V_max_ relative to DMSO but almost no change in K_M_. This background inhibition does not correlate with labeling and may be caused by nonspecific factors, such as aggregation of **2** at higher concentrations (≥ 50 µM). Data represent the mean of three independent assays ± the standard error of the mean. All assays were performed in the presence of 100 µM of β-mercaptoethanol.

To futher validate that **2** acts specifically through the K197 site and to explore the mechanism of inhibition by **2**, we solved a high-resolution (1.95 □, **Table 1**) crystal structure of K197C tethered with **2**. The structure confirms that **2** tethers to K197C rather than the active-site catalytic Cys215, and that tethered **2** resides in the 197 site (**Figure 8A**) rather than the relatively nearby BB site, which is also theoretically within reach of the tethering linker on the other side of the α3 helix. We modeled **2** as partially populated and, indeed, the 83% refined occupancy in the crystal structure was very similar to the ~85% conjugation measured after tethering in solution prior to crystallization. **2** adopts a conformation in which the two rings are nearly coplanar. This interpretation is further validated by a polder map, in which both the ligand and bulk solvent are omitted [36] (**Figure 8 - Figure S1**). While coplanar biphenyl rings are typically believed to be disfavored due to steric clashes, it is possible that hydrogen bonding of D148 with the phenol combined with the electronegativity of the para-fluoro leads to delocalization of the rings’ electrons and promotes a coplanar conformation. Additionally, **2** packs against the hydrophobic floor, centered on Leu172, of the relatively shallow binding pocket in the 197 site. Trapping of coplanar biphenyl rings covalently attached to a protein has previously been reported [37].

**Figure 8:**
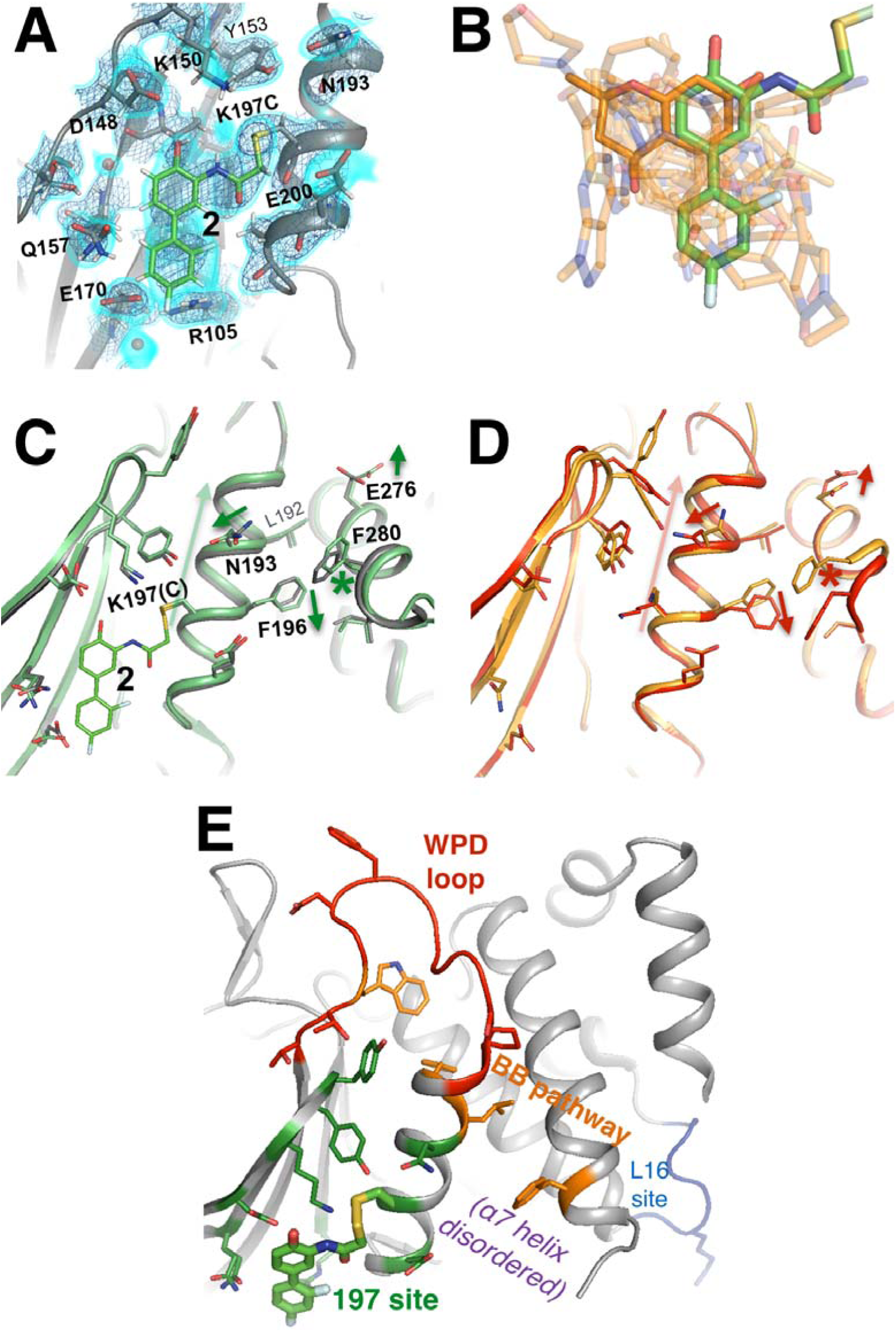
A functional small-molecule inhibitor tethered to the new 197 allosteric site. **(A)** The tethered inhibitor **2** is highly ordered (~85% occupancy) in the 197 site, as seen by 2Fo-Fc electron density for our 1.90 □ structure contoured at 0.75 σ (cyan volume) and at 1.5 σ (blue mesh) that is continuous to the K197C sidechain. A few waters (transparent red spheres) which appear to be mutually exclusive with the molecule are likely displaced by binding. **(B)** Many fragments from WT cocrystal structures (transparent orange) overlay well with **2** in the K197C cocrystal structure (green). One fragment in particular (solid orange) has a ring substructure that overlays very closely with a ring substructure of **2**. **(C)** The K197C **2**-tethered structure (green) is similar to the K197C apo structure (gray), but upon tethering there are several conformational changes (arrows and asterisk) in the α3 helix: the whole backbone shifts up in this view slightly leading back into the WPD loop (top), N193 switches rotamers, and the sidechains of F196 and E200 move within rotameric wells. The end of the α6 helix, including E276 and F280, appears to respond in concert. **(D)** Several of these changes mirror changes from open-to-closed apo PTP1B (arrows and asterisk) as seen in the two conformations of our 278 K model (red/orange). **(E)** Overview as in **Figure 1E** for context.

### Figure 8 - Movie S1

Movie version of **Figure 8A**.

**Figure 8 - Figure S1:**
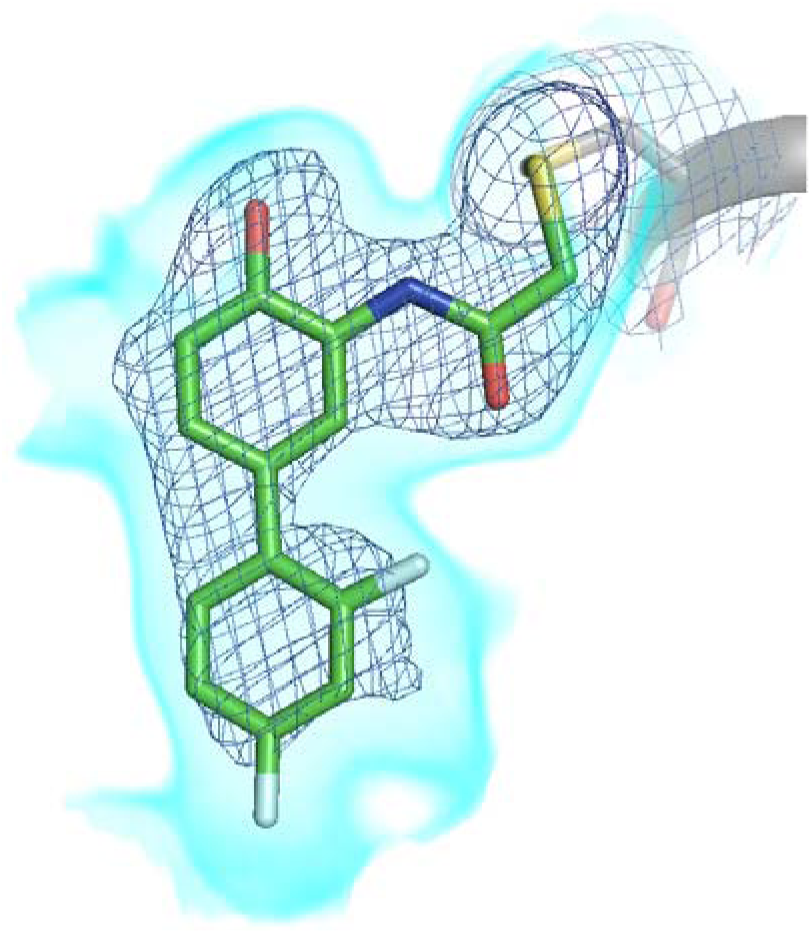
Polder omit map of K197C tethered structure. A polder map – an omit electron density map with bulk solvent excluded – is shown contoured at 1.5 σ (cyan volume) and at 3.5 σ (dark mesh). The ligand is shown here, but was not present during calculation of the map.

To elucidate confomational changes induced by **2**, we also solved a high-resolution (1.95 □, **Table 1**) crystal structure of apo K197C in the same crystal form for comparison. As mentioned previously, PTP1B remains in the open state without glycerol; glycerol was absent from the tethered K197C structure (to avoid competition for binding in the 197 site) and from the K197C apo structure (for consistency), so we are unable to see any dramatic shifts in the global open-closed equilibrium that **2** may induce. Tyr153 shifts its position slightly and Tyr152 responds by shifting fully to its up rotamer, but this is likely due to the loss of interactions with the WT K197 upon mutation. Beyond these mutation-induced effects, we see some conformational changes associated with tethering of **2**. The key residue Asn193 [11] changes rotamers, the sidechain of Phe196 on α3 “slides” to change its aromatic stacking arrangement with Phe280 on α6 (**Figure 8C**), and Glu276 – which contacts the wedge residue Leu192 [11] – rotates sidechain dihedral angles. These sidechain movements appear to couple to subtle, more distributed backbone shifts [38] of the α3 helix, several residues of which move up toward the WPD loop by ~0.5 □ (**Figure 8C**). Interestingly, these sidechain and backbone movements are somewhat similar to those between the two macrostates of apo PTP1B at high temperature (**Figure 8D**). Thus, although some mechanistic details remain unclear, allosteric inhibition by **2** may involve conformational changes, especially of α3, that are similar to those that occur during the global transition from the open to the closed state [11]. This interpretation is consistent with a recent report that mutations (Y153A, M282A) in what we here recognize as the 197 site and L16 site shift the WPD loop equilibrium toward an open, inactive state [24].

Several of the other noncovalent fragments bound to WT overlay well with the aromatic rings of **2** tethered to K197C (**Figure 8B**). This structural convergence suggests promising avenues for medicinal chemistry: moieties from fragments could be appended to **2** to create superior covalent inhibitors, or portions of **2** could be appended to fragments to create superior non-covalent inhibitors.

## Discussion

Our analysis of PTP1B paints a portrait of an inherently allosteric system. Allostery is fundamentally tied to protein functions such as catalysis via the theme of conformational motions [39]. Here we have harnessed new approaches in X-ray crystallography to map the coordinated conformational redistributions that underlie allostery in the dynamic enzyme PTP1B. Metaphorically, we were able to use this map of PTP1B’s “intramolecular nervous system” to target its allosteric “pressure points” and modulate its function.

Proteins sample many conformations from a complex energy landscape [40], many of which are accessible and represented among the millions to trillions of molecules in a protein crystal. However, an X-ray crystallographic dataset provides only ensemble-averaged information – so it is difficult to decipher individual minor conformations from a single dataset. A key to our work was harnessing the power of *en masse* structural analysis, which let us reveal minor conformations and the shifts between them that allow a dynamic protein to function. We exploited families of structures in two different ways. First, we contrasted structures at several different temperatures [12] for PTP1B to track coordinated conformational shifts which underlie allosteric communication. Second, we used hundreds of structures of PTP1B with different small-molecule fragments to calculate a statistical “background” electron density map representing the unbound state, which we could subtract to reveal fragment-bound conformations [14] for many allosteric sites. This requires using the PDB format of alternative locations to encode both compositional and conformational heterogeneity within a single model. Our multi-structure equilibrium X-ray approaches complement other methods for breaking the degeneracy of ensemble-averaged data to resolve multiple conformations of macromolecules. For example, 3D classification algorithms in cryo-electron microscopy enable *in silico* purification of different compositional and conformational states [41]. Time-resolved X-ray experiments, e.g. with free-electron lasers, now offer great promise for mapping conformational changes with both spatial and temporal resolution, although general experimental strategies are still forthcoming for the vast majority of proteins that are non-photoactivatable [42]. More generally, integrative modeling algorithms can synthesize data from disparate sources at different resolutions, including solution NMR or small-angle X-ray scattering, to build ensembles of structures that are consistent with all the experimental data [43,44].

Previous studies using the BB inhibitors [10] and mutations [11] revealed allosteric conformational changes linking distal areas to the active site which, presumably, were induced by the allosteric perturbations. In this work we demonstrate that, to a large degree, these alternative conformations are already latently sampled by the apo protein, and are simply further stabilized by the allosteric perturbations. Interestingly, the previously described allosteric regions are all situated on a contiguous face on the “back side” of the protein, centered around the quasi-ordered α7 helix (**Figure 1E, Figure 8E**). Here we have not only validated this collective allosteric network, but also identified major extensions of the network which extend in both directions from the classical BB allosteric pathway. Using covalent tethering, we have also investigated one such extension in more detail and confirmed that it is allosterically linked to enzyme function. Overall, our analysis suggests that allostery in PTP1B is characterized by a menagerie of inter-dependent conformational changes spanning length scales: from helical order-disorder transitions, to subtle helical tweaks, to local sidechain rotamer changes, to large discrete active-site loop transitions.

A central challenge in protein biophysics is coupling ligand binding to changes in the complex protein conformational ensemble that elicit a change in function. Our tethering screen of the 197 site in PTP1B (**Figure 7 - Figure S1**) suggests that although many small molecules can bind with appreciable occupancy to an allosteric site, most are benign – few inhibit activity upon binding like **2**. The large number of protein-ligand cocrystal structures we report here sets the stage to dissect this relationship between small-molecule binding and perturbations to specific alternative conformations at allosteric sites that are coupled to the active site, including the 197 site and L16 site. In the 197 site in particular, we have many structures with low-affinity, non-covalent small-molecule fragments (which led us to ligandable sites), and one structure with a covalently tethered molecule (which confirmed allosteric inhibition). Moving forward, pieces of non-covalent molecules could be appended to the covalent **2** to improve its inhibitory properties – but ultimately a covalent inhibitor must be redesigned to non-covalently target the wildtype protein. By contrast, pieces of the smaller non-covalent molecules, or of **2**, could be combined in many more ways to perhaps create a potent non-covalent inhibitor – but the combinatorial chemical space is more daunting. New higher-affinity molecules from either approach could improve allosteric inhibition by stabilizing specific conformations in the allosteric site associated with the closed WPD loop – or probably equally well by stabilizing specific conformations associated with the open WPD loop. Alternatively, they could leave the open-closed equilibrium unchanged, but modulate the interconversion rates [45] to interfere with catalysis. Future studies with new allosteric ligands of PTP1B at the 197 site or the L16 site will clarify the structural and dynamic mechanisms by which specific molecular species function as efficient allosteric inputs into protein receivers.

In addition to these newly available medicinal chemistry avenues, two features of the allosteric sites we have identified in PTP1B may be used in the future to strengthen binding affinity. First, some proteins contain reactive cysteines that are amenable to covalent ligands [46], but many others contain reactive lysines [47]. The 197 site in PTP1B of course contains K197, whose sidechain is roughly the same length as the tether arm linking **2** to the mutated K197C (**Figure 8**), suggesting that a targeted lysine-tethering strategy may be feasible in this case. A recent survey of lysine reactivity using a small set of ligands in >2,000 proteins, including PTP1B, did not identify K197 as highly reactive [47]; however, sampling more chemical space could reveal ligands that bind the 197 site via K197 specifically and tightly, as was the case for binding the 197 site via K197C in this study.

The second feature of the 197 site that could help improve allosteric inhibition is that, as stated above, the quasi-disordered α7 helix is capable of reordering into different conformations, some of which cover the α3-α6 region including the 197 site. We observe several reordered α7 conformations under different conditions: with BB3 at room temperature, with BB1 at cryogenic temperature, in the S295F (α7) mutant, in the L192A (α3) mutant with an active-site inhibitor (**Figure 2 - Figure S3C**), and with a fragment in the BB site (**Figure 6A**). Similarly, the disordered C-terminus, including α7, accommodates a recently reported allosteric inhibitor [7]. For the BB site, the malleability of α7 is likely a double-edged sword. On one hand, reordered conformations of α7 may contribute binding energy to allosteric inhibitors by forming a “lid” over what is partially a “cryptic site” [23]. On the other hand, they may also accommodate different small-molecule variants equally well, such that it is difficult to improve upon inhibition – i.e., the structure-activity relationship (SAR) is flat. However, the 197 site is structurally distinct from the BB site, and is accessible to different, more C-terminal portions of α7. It is therefore possible the 197 site will yield improved allosteric inhibitors that take advantage of these unique (quasi-)structural features.

Together, our observations suggest that the allosteric network we establish here within the catalytic domain of PTP1B may be a “receiver” for allosteric inputs from the C-terminus. If so, this strategy operates in parallel with other regulatory mechanisms such as active-site oxidation [31] and phosphorylation of other sites in the catalytic domain [48]. Interestingly, different protein tyrosine phosphatases (PTPs) feature a generally structurally conserved catalytic domain, but different variants of α7 or even entirely different N- or C-terminal domains [49], which can be trapped in inhibitory conformations for allosteric inhibition as recently realized for SHP2 [50]. Dissecting the ramifications of these customized molecular inputs for differential allosteric control of these PTPs to encode their unique cellular roles and disease phenotypes will be an exciting area of future research.

## Materials and Methods

### Cloning, expression, and purification

For “wildtype” PTP1B we used the 1-321 construct, with the C32S/C92V double mutation [51] to prevent off-target tethering reactions, in a pET24b vector with kanamycin resistance. K197C, K197A, Y152G, and Y153A were created using site-directed mutagenesis from that construct.

Protein was expressed and purified as previously reported [17], with some minor variations. For expression, we transformed BL21 E. coli cells with plasmid, grew cells on LB + kanamycin plates overnight at 37°C, inoculated 5 mL starter cultures of LB + kanamycin with individual colonies, grew shaking overnight at 37°C, inoculated larger 1 L cultures of LB + kanamycin, grew shaking at 37°C until optical density at 600 nm was approximately 0.6-0.8, induced with 100 mM IPTG, and grew shaking either for 4 hours at 37°C or overnight at 18°C. Cell pellets (“cellets”) were harvested by centrifugation and stored at -80°C in 50 mL conical tubes.

For purification, we first performed cation exchange with a SP FF 16/10 cation exchange column (GE Healthcare Life Sciences) in lysis buffer (100 mM MES pH 6.5, 1 mM EDTA, 1 mM DTT) with a multi-stage 0-1 M NaCl gradient (shallow at first for elution of PTP1B, then steeper); PTP1B eluted around 200 mM NaCl. We then performed size exclusion with a Superdex 75 size exclusion column (GE Healthcare Life Sciences) in size exclusion buffer (100 mM MES pH 6.5, 1 mM EDTA, 1 mM DTT, 200 mM NaCl). PTP1B appeared highly pure in SDS-PAGE gels.

### Crystallization

WT PTP1B was dialyzed into crystallization buffer (10 mM Tris pH 7.5, 0.2 mM EDTA, 25 mM NaCl, 3 mM DTT) with at least a 200x volume ratio overnight at 4°C. We were unable to grow apo WT PTP1B crystals initially, so we synthesized the active-site inhibitors OBA and OTP as in [52] (OBA = compound **3a**, OTP = compound **12h**). We were unable to solubilize OTP as used in [17]. Instead, we co-crystallized PTP1B with OBA [53]. We first solubilized OBA to 250 mM in DMSO, then created a 10:1 molar ratio of PTP1B:OBA. Crystallization drops were set in 96-well sitting- or hanging-drop format at 4°C with 10-15 mg/mL protein with 1 μL of protein solution + 1 μL of well solution (0.2-0.4 M magnesium acetate, 0.1 M HEPES pH 7.3-7.6, 12-17% PEG 8000), then trays were incubated at 4°C. Crystals several hundred μm long grew within a few days, and often continued to grow bigger for several more days. We created seed stocks from these crystals by pipetting the entire drop into 50μL of well solution, iterating between vortexing for 30 s and sitting on ice for 30 s several times, and performing serial 10-fold dilutions in well solution. Apo crystals were grown by introducing seed stock (0x, 10x, or 100x diluted) into freshly set drops, either by streaking with a cat whisker or pipetting a small amount (e.g. 0.1 μL into a 2 μL drop). Serial seeding using new apo crystals successively improved crystal quality. We also added ethanol to the well solution based on an additive screen (Hampton Research), and added glycerol to mimic the previously published apo structure protocol [17], resulting in the following final WT PTP1B crystallization well solution: 0.3 M magnesium acetate, 0.1 M HEPES pH 7.5, 0.1% β-mercaptoethanol, 16% PEG 8000, 2% ethanol, 10% glycerol. The resulting crystals were used for our WT PTP1B multitemperature analysis.

We also crystallized WT PTP1B in MRC SwissCi 3-well sitting-drop trays. Protein was at 30-50 mg/mL protein in the same crystallization buffer. The well solution was very similar except for having a slightly lower precipitant concentration (13-14% PEG 8000) and no glycerol. Drops were set at room temperature with 135 nL protein solution + 135 nL well solution + 30 nL seed stock, then trays were incubated at 4°C. Crystals appeared within a few days. The best seed stocks had been diluted 10-100x. These crystals were used for the BB3-soaking and fragment-soaking experiments.

We crystallized apo K197C in the microbatch format with Al’s oil covering all wells. Protein was at 5-30 mg/mL in the same crystallization buffer as WT but without DTT. The well solution was 0.3 M magnesium acetate, 0.1 M HEPES pH 7.5, 0.1% β-mercaptoethanol, 10-26% PEG 8000, 2% ethanol. Drops were set on ice with 1 μL protein solution + 1 μL well solution, then trays were incubated at 4°C. Crystals appeared within a few days. We also grew apo K197C crystals in a few other similar conditions.

We crystallized K197C tethered to **2** in the 96-well hanging-drop format. Protein was at 15 mg/mL in the same crystallization buffer as WT but without DTT. The well solution was 0.2 M magnesium acetate tetrahydrate, 20% PEG 3350. Drops were set at room temperature with 100 nL protein solution + 100 nL well solution, then trays were incubated at 4°C. Crystals appeared within a few days.

### X-ray data collection

We used PDB ID 1sug for the apo WT 100 K dataset. The apo WT 278 K dataset was collected at Stanford Synchrotron Radiation Lightsource (SSRL) beamline 12-2. All fragment-soaked datasets were collected at Diamond Light Source beamline I04-1. All other datasets were collected at Advanced Light Source (ALS) beamline 8.3.1.

For apo WT 180, 240, and 278 K, crystals had been grown in 2 μL with 10% glycerol in the mother liquor, then 1.5 μL of 50% glycerol was added several hours before data collection, resulting in a final concentration of ~27% glycerol. Some crystals were also dabbed into more 50% glycerol just before mounting.

For BB3-complexed WT, crystals were soaked with 125 nL of 10 mM BB3 (in DMSO). No glycerol was present in these crystals.

For apo WT and BB3-complexed WT, crystals were looped and placed in a plastic capillary with ~70% mother liquor, ~30% water to prevent dehydration during data collection, regardless of temperature; datasets were obtained at different temperatures simply by adjusting the cryojet temperature before placing the crystal on the goniometer. Helical data collection (translation along the crystal coupled to goniometer rotations) was used to expose fresh regions with each shot, to minimize radiation damage.

For apo and **2**-tethered K197C, crystals were simply looped and directly mounted on the goniometer in front of the cryojet. For apo K197C, a small amount of ice was likely present on the crystal.

For fragment-soaked PTP1B, WT PTP1B crystals in MRC SwissCi 3-well sitting-drop trays were soaked with small-molecule fragments using acoustic droplet ejection technology and a database mapping individual fragments to individual crystals as described [13]. PTP1B crystals were quite tolerant to DMSO, so we were able to achieve high fragment concentrations and long incubation times: we soaked overnight for >8 hr at final concentrations of 30% DMSO and 30-150 mM fragment (depending on the concentration of the fragments in the source library). Additionally we collected X-ray data for 48 “apo” datasets (soaked with DMSO), 42 of which gave high-resolution datasets, to better establish the unbound background electron density for PanDDA analysis. Despite the high DMSO concentrations, we did not observe difference electron density consistent with any ordered DMSO molecules bound to PTP1B. Some fragments were soaked into additional crystals if good datasets were not obtained from the initial soak; however, only 2 of the 110 fragment-bound datasets contain the same fragment. We also collected a relatively small number of trial datasets (28) at room temperature instead of cryogenic temperature, but they were generally low-resolution, and none revealed bound fragments.

Most crystals stuck to the bottoms of wells regardless of construct and tray format, but it was often possible to gently dislodge them, or to physically break them off then expose the unperturbed portion of the crystal to the X-ray beam. Each dataset in this study was collected from a single crystal.

### X-ray data processing

To process the multitemperature and tethered datasets, we used XDS [54]. In each case we chose a resolution cutoff for which CC1/2 [55] was statistically significant at the 0.1% level (above 0.4). We created a new set of R_free_ flags for the 278 K WT apo dataset hiGOL 278, then transferred them to the MTZ file of every other dataset with the reflection file editor in PHENIX [56] (for PDB ID 1sug, we first deleted the existing R_free_ flags). We solved each structure by molecular replacement with Phaser [57]. One solution was obtained for each dataset. For WT, we used PDB ID 1c83 with all waters and the WPD loop removed. For K197C, we used a refined WT PTP1B model for molecular replacement.

For fragment-soaked datasets, we used XDS and custom scripts to automatically determine resolution cutoffs for all datasets. The resolution cutoff was initialized at 1.4 □ and incremented until the following criteria were met for the highest-resolution bin: at least 1.0 I/σ(I), at least 50% CC1/2 [55], and at least 90% completeness. R_free_ flags were created for the highest-resolution dataset by transferring and extending the flags from the 278 K WT apo dataset using the Phenix reflection file editor. These R_free_ flags were then transferred from that highest-resolution dataset to every other dataset in the fragment-soaking experiment. For PanDDA to accept the MTZ files as inputs, it was necessary to modify each file so that all columns (H, K, L, structure factors, map coefficients, and R-free flags) had the same number of indices; no observations were omitted in this step. We then phased each dataset with Phaser using a reference model that was created by interpreting a high-resolution DMSO-soaked apo dataset. Next, we refined each initial model from Phaser using phenix.refine with the follwing flags to prevent excessive coordinate drift: “reference_coordinate_restraints.enabled=True “reference_coordinate_restraints.sigma=0.1”. Structure factors were then dropped from MTZ files, leaving map coefficients as inputs to PanDDA. Filled map coefficients (from Phenix) were used to avoid Fourier series truncation effects in PanDDA maps. The resulting models were used as input to PanDDA (see below).

### Structure modeling

For **Figure 1 - Figure S1**, we re-refined the following 36 structures of PTP1B from the PDB either as-is, or with a dual-conformation WPD loop: 1bzj 1kak 1oem 1oeo 1sug 1t48 1t49 1t4j 2azr 2b07 2bgd 2f6f 2f6t 2f6v 2f6w 2f6y 2f6z 2f70 2f71 2h4k 2hb1 2qbp 2qbq 2qbr 2qbs 2zmm 2zn7 3cwe 3d9c 3eax 3eb1 3eu0 3i7z 3i80 3sme 4i8n. For the dual-conformation refinements, we constrained occupancies of the open + closed conformations of the WPD loop to 1.

For multitemperature WT, we refined the initial model using phenix.refine for 10 macrocycles with automated water picking turned off. Next, we inserted preliminary open and closed alternative conformations for the WPD loop, and refined for another 10 macrocycles with automated water picking turned on. Finally, we performed several rounds of manual rebuilding, including manual addition and deletion of protein, solvent, and glycerol conformations and refinement with automated water picking turned off. Anisotropic B-factors were not used in refinement. The α7 helix was modeled as alternative conformation A only, with the unmodeled B conformation presumed to correspond to the disordered state; this allowed the occupancy of the ordered state to be refined. It was necessary to provide explicit occupancy parameter files to phenix.refine in some cases. For many residues, conformations obtained from PDB ID 1sug or 1t49 or another of our datasets (usually higher temperatures) were useful for “filling in” missing density. Often the missing conformations would not have been obvious based on the map alone, but once inserted and refined they seemed to fit well. This cross-dataset conformational-sampling approach also had the effect of emphasizing differences between models from different temperatures while minimizing differences due to chance or arbitrary modeling choices. Nevertheless, we encourage future users of these datasets to compare across different temperatures based at least in part on the electron density, and not just our models. The building process was guided by all-atom structure validation with MolProbity [58]. The 100 K WT model (1sug) is truly WT, whereas our new “WT” datasets are all C32S/C92V; however, both cysteine-scrubbing mutations are structurally conservative, distal to the active site, and apparently uncoupled from the WPD loop and all allosteric regions.

Glycerol (~20% final) was present in WT crystals during each multitemperature data collection to maintain consistency with the 100 K structure (PDB ID 1sug), in which glycerol was used as a cryoprotectant. Several ordered glycerol molecules, including those in contact with the closed WPD loop and at the 197 allosteric site, were evident from electron density at all temperatures. However, in some cases it was difficult to differentiate between ordered waters, glycerols, or simply noise in the map. For example, the electron density was uncertain at some of the elevated temperatures for some glycerols originally modeled in PDB ID 1sug. When glycerol was omitted from crystals, the WPD loop was entirely in the open conformation regardless of temperature, from cryogenic to room-temperature (data not shown). Our interpretation is that ordered glycerols in the active site, which are evident from the electron density at all temperatures, make weak contacts with the WPD loop’s closed conformation, and thus shift the protein’s energy landscape to a regime in which the open vs. closed conformations are near enough to isoenergetic that temperature can modulate their populations. This interpretation is strengthened by the fact that these glycerol molecules align well with a bound mimic of the pTyr substrate (PDB ID 1pty), which causes the loop to close during the catalytic cycle.

For K197C, we used a similar refinement procedure, including many manual tweaks of alternative conformations for protein and water atoms. For the **2**-tethered K197C structure, we omitted **2** for many rounds of refinement, allowing the electron density for the missing molecule to become extremely convincing before we finally added it to the model. The distance between the sulfur atoms in K197C and the ligand was restrained to 2.15 with a σ of 0.1 for refinement.

For fragment-soaked datasets, we used the PanDDA approach [14] in a few stages. First, using PanDDA version 0.1, we ran pandda.analyse, and interpreted and modeled events. Next, using the new PanDDA version 0.2, we ran pandda.analyse again. During this second run, datasets which were events in the first run were excluded from background density calculation, and datasets that had substantial map artifacts or very noisy/low-quality maps in the first run were excluded entirely.

Some events for which we modeled bound fragments in the earlier PanDDA version 0.1 runs were not detected as events in the final PanDDA version 0.2 run. In these cases, we manually created event maps based on the 1-BDC background subtraction values from the earlier PanDDA runs. In most cases, visual inspection confirmed that these early events likely correspond to bound fragments.

Many PanDDA “events” in the active site corresponded not to ligand binding, but rather to protein conformational changes of the WPD loop, P-loop, and Y46 loop that are caused by oxidation of the catalytic Cys215, a natural PTP1B regulatory mechanism [31]. Some other active-site events were difficult to interpret, perhaps due to active-site dynamics or differences in the appropriate background model for the open vs. closed state of the protein; future methodological improvements may clarify modeling in such cases.

We built a generic unbound-state model by interpreting both an average map for the highest-resolution bin and one of the best individual apo datasets. The WPD loop was modeled as open, Tyr152 was modeled with two alternative sidechain conformations on the loop 11 backbone that is compatible with the open WPD loop, and the N-terminus (start of α1) and C-terminus (end of α6, since α7 was disordered) were fit as well as possible. Ordered waters were also manually positioned. This generic unbound-state model was superposed onto each PanDDA input model in the correct reference frame, then refined, to create an unbound-state model for each dataset.

For each fragment-bound state, we inspected the fragment binding site, plus the several interesting regions of the protein mentioned above, in detail interactively. Waters were copied over from the unbound-state model, then moved or deleted where they conflicted with the bound fragment and/or the PanDDA event map. For a small number of planar fragments, several copies of the fragment bind in a parallel stack bridging the 197 site and a symmetry-related copy of the WPD-loop residue Phe182 via a crystal-lattice contact. Some areas such as Tyr152 were modeled with alternative conformations in the bound state only when they were well justified in the event map; otherwise we generally adhered to the unbound-state model. The correct modeling choice for the termini was uncertain in some cases.

To refine structures for the 110 datasets with one or more modeled fragments, first we created restraints files for the ligands with eLBOW [59]. For a small number of ligands, we additionally used AceDRG [60] and found that AceDRG generated more realistics restraints. Next, the pandda.export method in PanDDA version 0.2 was used to create an “ensemble structure” containing both the unbound state (including alternative conformations) and the bound state (including alternative conformations) in one multiconformer model. In pandda.export, the parameter “options.prune_duplicates_rms=0.2” was used to merge alternative conformations that were highly similar, and the parameter “duplicates.rmsd_cutoff=0.4” was used to restrain the coordinates of somewhat similar alternative conformations to be identical. These parameter values were chosen to effectively merge residues with very similar coordinates, while still allowing residues we evaluated as having genuine alternative conformations to remain separate and unrestrained. The resulting geometry restraint files from pandda.export are necessary to minimize overfitting or coordinate drift during refinement of this model type.

For refinement of the ensembles representing multiconformer models of the apo and bound states, we first refined each ensemble structure with phenix.refine to obtain water positions. The first stage of Phenix refinement was 10 macrocycles with no removal or addition of waters (“ordered_solvent=False”) to let the existing waters relax into local minima. The second stage of Phenix refinement was another 10 macrocycles with automated removal and addition of waters (“ordered_solvent=True”) to remove waters that were unable to reach local minima and add waters that were clearly missing. Adding and removing waters, when compared to only removing them, generally had negligible effect on MolProbity scores, but improved R_work_ and R_free_. During this first Phenix refinement stage to obtain water positions, occupancies were fixed to the original PanDDA BDC value for the ground state and 1-BDC for the bound state; occupancy was distributed evenly between substates when the ground state or the bound state had alternative conformations for some residues. We observed coordinate drift and unstable B-factors for the protein with Phenix refinement. Therefore, we copied the water positions obtained from Phenix into the initial ensemble models, and refined with Refmac [61]. To do so, we first set all B-factors to 40 □^2^, set bound-state occupancies to 2*(1-BDC) and unbound-state occupancies to (with occupancy evenly distributed across alternative conformations within each state), and generated new restraints files that included the water molecules by running the PanDDA utility giant.make_restraints with the same RMSD parameter as with giant.merge_conformations: “duplicates.rmsd_cutoff=0.4”. Then, each ensemble model was refined with Refmac using giant.quick_refine with the ligand cif and custom giant.make_restraints restraint parameter files using the protocol herein. First, each model was refined for the default 10 cycles, with the extra arguments to Refmac "MAKE HOUT Yes", to preserve hydrogens, and “HOLD 0.001 100 100" to restrain XYZ coordinates but still allow for some geometry regularization and encourage B-factor and occupancy convergence. Next, the output from that refinement was fed into a loop of Refmac refinement with the default 10 cycles per run, and the “HOLD 0.0001 100 100” argument, essentially fixing the XYZ coordinates, while letting occupancies and B-factors refine. Our output was the result of the 4th round (1 round + 3 rounds) of refinement in Refmac. However, the occupancies refined with these refinements did not converge to the correct occupancy (as seen by huge difference peaks describing the ligand). We then refined these structures in phenix, fixing the XYZ coordinates and manually scanning across possible occupancies while refining B-factors with the following settings: refinement.refine.strategy = individual_adp, hydrogens.optimize_scattering_contribution = False, main.number_of_macro_cycles = 10, optimize_mask = True, optimize_adp_weight = True. While in principle one could interpret the difference density to pick an optimal refined occupancy, no other statistics calculated provided a clear choice of occupancy. We ultimately chose to deposit occupancies of the bound state at 2.2 times the event map occupancy (1-BDC). This occupancy choice was motivated by the trend previously found by Pearce et al [14]. In cases where the total bound occupancy was 50% or higher, the models were manually inspected, and a few dropped to low occupancies that minimized difference features of the ligand. The resulting final ensemble structures of the unbound state plus the fragment-bound state were converted from PDB to mmCIF format and deposited in the PDB using the new multimodel submission procedure.

### Visualization

Coot [62] was instrumental to visualizing and interactively modeling all structures. PyMOL [63] was used for all molecular graphics after initial modeling. We frequently used the volume rendering feature for low-contour electron density alongside the traditional mesh for higher-contour electron density.

### Synthesis of tethered compounds

#### Synthesis of 3-amino-2',4'-difluoro-[1,1'-biphenyl]-4-ol

A solution of 2,4-difluorophenyl boronic acid (188 mg, 1 mmol, 1.0 equiv), 2-amino-4-bromophenol (7.2 mg, 0.62 mmol, 1.2 equiv), Pd(PPh_3_)_4_ (57.8 mg, 0.05 mmol, 0.05 equiv), and Na_2_CO_3_ (318 mg, 3 mmol, 3 equiv) in THF (8 mL) was stirred at 90 ˚C overnight. The reaction was allowed to cool, taken up in water, and then extracted 3x with EtoAC. Organic layers were combined, washed with brine, dried over Na_2_SO_4_, concentrated *in vacuo* and purified using flash chromatography (MeOH/DCM) to obtain 22.1 mg of product (10% yield). Calcd for C_12_H_10_F_2_NO (M+H^+^): 222.07; Found 222.8.

#### Synthesis of **2**

To a mixture of 3-amino-2',4'-difluoro-[1,1'-biphenyl]-4-ol (22.1 mg, 0.1 mmol, 1 equiv), dithiodiglycolic acid (9.1 mg, 0.05 mmol, 0.5 equiv), HOBt·H_2_O (19.9 mg, 0.13 mmol, 1.3 equiv), and DIPEA (0.226 µL, 1.3 mmol, 1.3 equiv) in THF (300 µL), EDCI·HCl (25 mg, 0.13 mmol, 1.3 equiv) was added. The reaction was stirred at room temperature overnight. To this was added a solution of bis[2-(N,N-dimethylamino)ethyl]disulfide dihydrochloride (70.3 mg, 0.25 mmol, 2.5 equiv) and TCEP (2.5 mg, 0.01 mmol, 0.01 equiv) in THF/H_2_O (8:3, 275 µL). The reaction was stirred at room temperature overnight. The reaction was taken up in EtoAC/H_2_O, and extracted 3 times with EtOAc. Organic layers were combined, washed with brine, dried over Na_2_SO_4_, concentrated in vacuo and purified using flash chromatography (DCM/MeOH) followed by preparative HPLC (C_18_ column (50 x 19 mm), Methanol/Water–0.05% formic acid gradient: 10:90 to 100:0 over 12 min; 20 mL/min; 254 nm detection for 18 min.) to obtain 9.1 mg of **2** (23% yield). ^1^H NMR (400 MHz, Acetone-*d*_6_) δ 9.55 (s, 1H), 8.17 (s, 1H), 8.06-8.04 (m, 1H), 7.54-7.45 (m, 1H), 7.22-7.19 (m, 1H), 7.12-7.00 (m, 3H), 3.94-3.78 (m, 2H), 3.15-2.99 (m, 4H), 2.54-2.52 (m, 6H). Calcd for C_18_H_21_F_2_N_2_O_2_S_2_ (M+H^+^): 399.1; Found 398.93.

### Tethering

We screened K197C against a previously synthesized library of 1600 disulfide fragments made available by the UCSF Small Molecule Discovery Center (SMDC) [34,35].

For the screen, tethering reactions were performed using the following conditions: 1x tethering buffer (25 mM Tris pH 7.5, 100 mM NaCl), with 500 nM of K197C, 1 mM β-mercaptoethanol, and 100 µM of fragment (0.2% DMSO), 1 hr at rt. Unless otherwise noted, tethering reactions for follow up experiments and activity assays were performed using the following conditions: 1x tethering buffer, 1 µM of K197C, 0.1 mM β-mercaptoethanol, and 50 µM of fragment (2% DMSO), 1 hr at rt. For crystallography, tethering reactions were performed using the following conditions: 1x tethering buffer, 0.76 mg/mL of K197C, 0.1 mM β-mercaptoethanol, 500 µM of TCS401, and 250 µM of fragment (2% DMSO), 2 hr at rt. A total reaction size of 3.5 mL was used for preparation of crystallography samples. Following labeling, the reaction was dialyzed into crystallization buffer overnight. In all cases the percent of tethering was measured using a Waters Xevo G2-XS Mass Spectrometer, and calculated by comparing the relative peak heights of the unmodified and modified protein. Tethering EC_50_ values were calculated using nonlinear fitting in Prism 7 (Graphpad), n = 3.

### Activity assays

For activity assays of WT PTP1B vs. allosteric mutants (**Figure 4 - Figure S3**), protein was diluted to 215 nM (WT) or 200 nM (mutants) in a variant of pNPP activity assay buffer (50 mM HEPES pH 7, 100 mM NaCl, 1 mM EDTA, and 1 mM DTT). WT assays were performed at 215 nM protein and mutant assays were performed at 200 nM, so WT data is normalized to 200 nM in both panels in **Figure 4 - Figure S3**. Enzyme activity assays were performed across 10 p-nitrophenyl phosphate (pNPP) concentrations obtained by serial 2-fold dilutions starting from 20 mM. A no-enzyme well was also assayed. Absorbance at 405 nm for each reaction was monitored every 30 s for 5 minutes using a Tecan Infinite M200 Pro. The rate (mAu/s) of each reaction was calculated over the 5 minutes. Michaelis-Menten parameters were then calculated using Prism 7 (Graphpad). These parameters for WT PTP1B were similar to those reported previously [10].

For activity inhibition assays of WT PTP1B with small-molecule fragments, 20 fragments were chosen early in the iterative PanDDA analysis process (see “Structure modeling”). Protein was diluted to 200 nM in a variant of pNPP activity assay buffer (50 mM HEPES pH 7, 100 mM NaCl, 1 mM EDTA, 0.05% Tween-20, and 100 mM β-mercaptoethanol). Enzyme activity assays were performed with 0.15 or 1 mM fragment in 2% DMSO (final) or with 2% DMSO without fragment as a control, with 5 mM pNPP. A no-enzyme well was also assayed. Absorbance at 405 nm for each reaction was monitored every 30 s for 5 minutes. The rate (mAu/s) of each reaction was calculated over the 5 minutes. These rates were compared with fragment vs. with DMSO.

For single-point assays of tethered K197C, completed tethering reactions (post 1 hr incubation) were diluted to a final concentration of 200 nM K197C with a variant of pNPP activity assay buffer (50 mM HEPES pH 7, 100 mM NaCl, 1 mM EDTA, 0.05% Tween-20, and 100 mM β-mercaptoethanol) and pNPP (5 mM final). A no-enzyme well and a DMSO-only well were also assayed. Absorbance at 405 for each reaction was monitored every 30 s for 5 minutes using a Tecan Infinite M200 Pro. Percent inhibition was calculated using the following equation: 100(1- ((RateFragment-RateNo Enzyme)/(Rate_DMSO_-RateNo Enzyme)))

For **2** titration assays of tethered K197C and WT, completed tethering reactions (post 1 hr incubation) were were diluted to a final concentration of 100 nM K197C and WT with a variant of pNPP activity assay buffer (50 mM HEPES pH 7, 100 mM NaCl, 1 mM EDTA, 0.05% Tween-20, and 100 mM β-mercaptoethanol), 9 different concentrations of **2** obtained by serial 3-fold dilutions starting at 50 µM, and pNPP (5 mM final). A no-enzyme well and a DMSO-only well were also assayed. Absorbance at 405 nm for each reaction was monitored every 30 s for 5 minutes using a Tecan Infinite M200 Pro. The rate (mAu/s) of each reaction was calculated over the 5 minutes. Percent inhibition was calculated using the following equation: 100(1- ((RateFragment-RateNo Enzyme)/(Rate_DMSO_-RateNo Enzyme))). IC_50_ values were calculated using nonlinear fitting in Prism 7 (Graphpad), n = 3.

For kinetics experiments with tethered complexes, completed K197C and WT tethering reactions (post 1 hr incubation) were were diluted to a final concentration of 200 nM K197C and WT with a variant of pNPP activity assay buffer (50 mM HEPES pH 7, 100 mM NaCl, 1 mM EDTA, 0.05% Tween-20, and 100 mM β-mercaptoethanol), **2** (50 µM), and 10 pNPP concentrations obtained by serial 2-fold dilutions starting from 20 mM. Absorbance at 405 nm for each reaction was monitored every 30 s for 5 minutes using a Tecan Infinite M200 Pro. The rate (mAu/s) of each reaction was calculated over the 5 minutes. Data was plotted and fit using Prism 7 (Graphpad), n = 3.

### Data availability

We have also made publically available several files that document our PanDDA analysis of all WT fragment-soaked datasets. For each dataset, we provide a model of the unbound state, structure factors, an average map for the corresponding resolution bin, a PanDDA Z-map, and one or more PanDDA event map(s) as applicable. For fragment-bound datasets, we also provide the refined ground state model and the bound state model (before they were merged into an ensemble and refined) as separate PDB files, along with Phenix, Refmac, and ligand restraint files used in the ensemble refinement. Finally, we provide an overall PanDDA log file. These files are hosted at Zenodo at the following DOI: 10.5281/zenodo.1044103.

Multiconformer models and structure factors for the multitemperature WT (6B90, 6B8E, 6B8T, 6B8X), BB3-bound (6B8Z), K197C apo (6BAI) and tethered (6B95) datasets have been deposited in the Protein Data Bank [22]. Refined ensemble structures for the 110 WT fragment-bound datasets have been deposited using a new procedure and PDB IDs are pending (all data underlying data are accessible at Zenodo).

## Acknowledgements

We thank Michelle Arkin for helpful suggestions; the Small Molecule Discovery Center at UCSF for use of the disulfide-fragment library; Chris Wilson, Ken Hallenbeck, and Gregory Lee for assistance while performing the Tethering screen; Nicholas Rettko for help generating mutant PTP1B plasmids; Nigel Moriarty and Brandi Hudson for help with covalent ligand restraints; Patrick Collins, Alice Douangamath, and Tobias Krojer for help operating the XChem fragment-screening pipeline; and Anil Verma for operation of the Research Center at Harwell crystallization facility. We also “thank” LK for “help” with data collection.

J.S.F. is supported by a Searle Scholar Award from the Kinship Foundation, a Pew Scholar Award from the Pew Charitable Trusts, a Packard Fellowship from the David and Lucile Packard Foundation, NIH GM110580, UC Office of the President Laboratory Fees Research Program LFR-17-476732, and NSF STC-1231306.

J.A.W. is supported by NIH CA191018.

Z.B.H. is supported by a postdoctoral fellowship from the Helen Hay Whitney Foundation and HHMI, as well as a Pathway to Independence Award from the NIH-NCI (K99CA203002)

D.A.K. is supported by an A.P. Giannini Postdoctoral Fellowship.

J.T.B. is supported by an NSF Graduate Research Fellowship.

T.J.R. is supported by an NIH Predoctoral Fellowship (F31 CA180378-01).

Data collection at Beamline 8.3.1 at the Advanced Light Sources is is supported by the University of California Office of the President, Multicampus Research Programs and Initiatives grant MR-15-328599, the Program Breakthrough Biomedical Research (which is partially funded by the Sandler Foundation), the National Institutes of Health (R01 GM124149 and P30 GM124169), Plexxikon Inc., and the Integrated Diffraction Analysis Technologies program of the US Department of Energy Office of Biological and Environmental Research. The Advanced Light Source (Berkeley, CA) is a national user facility operated by Lawrence Berkeley National Laboratory on behalf of the US Department of Energy under contract number DE-AC02-05CH11231, Office of Basic Energy Sciences.

Use of the Stanford Synchrotron Radiation Lightsource, SLAC National Accelerator Laboratory, is supported by the U.S. Department of Energy, Office of Science, Office of Basic Energy Sciences under Contract No. DE-AC02-76SF00515. The SSRL Structural Molecular Biology Program is supported by the DOE Office of Biological and Environmental Research, and by the National Institutes of Health, National Institute of General Medical Sciences, P41GM103393.

## References

1. DeDecker BS. Allosteric drugs: thinking outside the active-site box. Chem Biol. 2000;7: R103–7. Available: https://www.ncbi.nlm.nih.gov/pubmed/10801477

2. Hardy JA, Wells JA. Searching for new allosteric sites in enzymes. Curr Opin Struct Biol. 2004;14: 706–715. doi:10.1016/j.sbi.2004.10.009

3. Zhang ZY. Protein tyrosine phosphatases: prospects for therapeutics. Curr Opin Chem Biol. 2001;5: 416–423. Available: https://www.ncbi.nlm.nih.gov/pubmed/11470605

4. Motlagh HN, Wrabl JO, Li J, Hilser VJ. The ensemble nature of allostery. Nature. 2014;508: 331–339. doi:10.1038/nature13001

5. Gunasekaran K, Ma B, Nussinov R. Is allostery an intrinsic property of all dynamic proteins? Proteins. 2004;57: 433–443. doi:10.1002/prot.20232

6. Elchebly M, Payette P, Michaliszyn E, Cromlish W, Collins S, Loy AL, et al. Increased insulin sensitivity and obesity resistance in mice lacking the protein tyrosine phosphatase-1B gene. Science. 1999;283: 1544–1548. Available: https://www.ncbi.nlm.nih.gov/pubmed/10066179

7. Krishnan N, Navasona K, Dorothy K, Miller DH, Bin X, Akshinthala SD, et al. Targeting the disordered C terminus of PTP1B with an allosteric inhibitor. Nat Chem Biol. 2014;10: 558–566. doi:10.1038/nchembio.1528

8. Krishnan N, Krishnan K, Connors CR, Choy MS, Page R, Peti W, et al. PTP1B inhibition suggests a therapeutic strategy for Rett syndrome. J Clin Invest. 2015;125: 3163–3177. doi:10.1172/JCI80323

9. Zhang Z-Y. Drugging the Undruggable: Therapeutic Potential of Targeting Protein Tyrosine Phosphatases. Acc Chem Res. 2017;50: 122–129. doi:10.1021/acs.accounts.6b00537

10. Wiesmann C, Barr KJ, Kung J, Zhu J, Erlanson DA, Shen W, et al. Allosteric inhibition of protein tyrosine phosphatase 1B. Nat Struct Mol Biol. 2004;11: 730–737. doi:10.1038/nsmb803

11. Choy MS, Li Y, Machado LESF, Kunze MBA, Connors CR, Wei X, et al. Conformational Rigidity and Protein Dynamics at Distinct Timescales Regulate PTP1B Activity and Allostery. Mol Cell. 2017;65: 644–658.e5. doi:10.1016/j.molcel.2017.01.014

12. Keedy DA, Kenner LR, Warkentin M, Woldeyes RA, Hopkins JB, Thompson MC, et al. Mapping the conformational landscape of a dynamic enzyme by multitemperature and XFEL crystallography. Elife. 2015;4. doi:10.7554/eLife.07574

13. Collins PM, Ng JT, Talon R, Nekrosiute K, Krojer T, Douangamath A, et al. Gentle, fast and effective crystal soaking by acoustic dispensing [Internet]. 2016. doi:10.1101/085712

14. Pearce NM, Krojer T, Bradley AR, Collins P, Nowak RP, Talon R, et al. A multi-crystal method for extracting obscured crystallographic states from conventionally uninterpretable electron density. Nat Commun. 2017;8: 15123. doi:10.1038/ncomms15123

15. Erlanson DA, Wells JA, Braisted AC. Tethering: fragment-based drug discovery. Annu Rev Biophys Biomol Struct. 2004;33: 199–223. doi:10.1146/annurev.biophys.33.110502.140409

16. Whittier SK, Hengge AC, Loria JP. Conformational motions regulate phosphoryl transfer in related protein tyrosine phosphatases. Science. 2013;341: 899–903. doi:10.1126/science.1241735

17. Pedersen AK, Peters G GÜH, Møller KB, Iversen LF, Kastrup JS. Water-molecule network and active-site flexibility of apo protein tyrosine phosphatase 1B. Acta Crystallogr D Biol Crystallogr. 2004;60: 1527–1534. doi:10.1107/S0907444904015094

18. Lang PT, Ng H-L, Fraser JS, Corn JE, Echols N, Sales M, et al. Automated electron-density sampling reveals widespread conformational polymorphism in proteins. Protein Sci. 2010;19: 1420–1431. doi:10.1002/pro.423

19. Fraser JS, van den Bedem H, Samelson AJ, Lang PT, Holton JM, Echols N, et al. Accessing protein conformational ensembles using room-temperature X-ray crystallography. Proceedings of the National Academy of Sciences. 2011;108: 16247–16252. doi:10.1073/pnas.1111325108

20. Diederichs K. Some aspects of quantitative analysis and correction of radiation damage. Acta Crystallogr D Biol Crystallogr. 2006;62: 96–101. doi:10.1107/S0907444905031537

21. Keedy DA, Fraser JS, van den Bedem H. Exposing Hidden Alternative Backbone Conformations in X-ray Crystallography Using qFit. PLoS Comput Biol. 2015;11: e1004507. doi:10.1371/journal.pcbi.1004507

22. Berman HM, Westbrook J, Feng Z, Gilliland G, Bhat TN, Weissig H, et al. The Protein Data Bank. Nucleic Acids Res. 2000;28: 235–242. Available: https://www.ncbi.nlm.nih.gov/pubmed/10592235

23. Cimermancic P, Weinkam P, Rettenmaier TJ, Bichmann L, Keedy DA, Woldeyes RA, et al. CryptoSite: Expanding the Druggable Proteome by Characterization and Prediction of Cryptic Binding Sites. J Mol Biol. 2016;428: 709–719. doi:10.1016/j.jmb.2016.01.029

24. Cui DS, Beaumont V, Ginther PS, Lipchock JM, Loria JP. Leveraging Reciprocity to Identify and Characterize Unknown Allosteric Sites in Protein Tyrosine Phosphatases. J Mol Biol. 2017;429: 2360–2372. doi:10.1016/j.jmb.2017.06.009

25. Bandyopadhyay D, Kusari A, Kenner KA, Liu F, Chernoff J, Gustafson TA, et al. Protein-Tyrosine Phosphatase 1B Complexes with the Insulin Receptor in Vivo and Is Tyrosine-phosphorylated in the Presence of Insulin. J Biol Chem. 1997;272: 1639–1645. doi:10.1074/jbc.272.3.1639

26. Rhee J, Lilien J, Balsamo J. Essential tyrosine residues for interaction of the non-receptor protein-tyrosine phosphatase PTP1B with N-cadherin. J Biol Chem. 2001;276: 6640–6644. doi:10.1074/jbc.M007656200

27. Smith CA, Ban D, Pratihar S, Giller K, Schwiegk C, de Groot BL, et al. Population shuffling of protein conformations. Angew Chem Int Ed Engl. 2015;54: 207–210. doi:10.1002/anie.201408890

28. Lovell SC, Word JM, Richardson JS, Richardson DC. The penultimate rotamer library. Proteins. 2000;40: 389–408. Available: http://www.ncbi.nlm.nih.gov/pubmed/10861930

29. Murray CW, Blundell TL. Structural biology in fragment-based drug design. Curr Opin Struct Biol. 2010;20: 497–507. doi:10.1016/j.sbi.2010.04.003

30. Hartshorn MJ, Murray CW, Cleasby A, Frederickson M, Tickle IJ, Jhoti H. Fragment-based lead discovery using X-ray crystallography. J Med Chem. 2005;48: 403–413. doi:10.1021/jm0495778

31. van Montfort RLM, Congreve M, Tisi D, Carr R, Jhoti H. Oxidation state of the active-site cysteine in protein tyrosine phosphatase 1B. Nature. 2003;423: 773–777. doi:10.1038/nature01681

32. Mattos C, Ringe D. Locating and characterizing binding sites on proteins. Nat Biotechnol. 1996;14: 595–599. doi:10.1038/nbt0596-595

33. Erlanson DA, Braisted AC, Raphael DR, Randal M, Stroud RM, Gordon EM, et al. Site-directed ligand discovery. Proc Natl Acad Sci U S A. 2000;97: 9367–9372. Available: https://www.ncbi.nlm.nih.gov/pubmed/10944209

34. Kathman SG, Xu Z, Statsyuk AV. A fragment-based method to discover irreversible covalent inhibitors of cysteine proteases. J Med Chem. 2014;57: 4969–4974. doi:10.1021/jm500345q

35. Burlingame MA, Christopher T M, Renslo AR. Simple One-Pot Synthesis of Disulfide Fragments for Use in Disulfide-Exchange Screening. ACS Comb Sci. 2011;13: 205–208. doi:10.1021/co200038g

36. Liebschner D, Afonine PV, Moriarty NW, Poon BK, Sobolev OV, Terwilliger TC, et al. Polder maps: improving OMIT maps by excluding bulk solvent. Acta Crystallogr D Struct Biol. 2017;73: 148–157. doi:10.1107/S2059798316018210

37. Pearson AD, Mills JH, Song Y, Nasertorabi F, Han GW, Baker D, et al. Trapping a transition state in a computationally designed protein bottle. Science. 2015;347: 863–867. doi:10.1126/science.aaa2424

38. Deis LN, Pemble CW 4th, Qi Y, Hagarman A, Richardson DC, Richardson JS, et al. Multiscale conformational heterogeneity in staphylococcal protein a: possible determinant of functional plasticity. Structure. 2014;22: 1467–1477. doi:10.1016/j.str.2014.08.014

39. Goodey NM, Benkovic SJ. Allosteric regulation and catalysis emerge via a common route. Nat Chem Biol. 2008;4: 474–482. doi:10.1038/nchembio.98

40. Frauenfelder H, Sligar SG, Wolynes PG. The energy landscapes and motions of proteins. Science. 1991;254: 1598–1603. Available: https://www.ncbi.nlm.nih.gov/pubmed/1749933

41. Scheres SHW. Processing of Structurally Heterogeneous Cryo-EM Data in RELION. Methods Enzymol. 2016;579: 125–157. doi:10.1016/bs.mie.2016.04.012

42. Hekstra DR, Ian White K, Socolich MA, Henning RW, Šrajer V, Ranganathan R. Electric-field-stimulated protein mechanics. Nature. 2016;540: 400–405. doi:10.1038/nature20571

43. van den Bedem H, Fraser JS. Integrative, dynamic structural biology at atomic resolution–it’s about time. Nat Methods. 2015;12: 307–318. doi:10.1038/nmeth.3324

44. Russel D, Lasker K, Webb B, Velázquez-Muriel J, Tjioe E, Schneidman-Duhovny D, et al. Putting the pieces together: integrative modeling platform software for structure determination of macromolecular assemblies. PLoS Biol. 2012;10: e1001244. doi:10.1371/journal.pbio.1001244

45. Popovych N, Sun S, Ebright RH, Kalodimos CG. Dynamically driven protein allostery. Nat Struct Mol Biol. 2006;13: 831–838. doi:10.1038/nsmb1132

46. Backus KM, Correia BE, Lum KM, Forli S, Horning BD, González-Páez GE, et al. Proteome-wide covalent ligand discovery in native biological systems. Nature. 2016;534: 570–574. doi:10.1038/nature18002

47. Hacker SM, Backus KM, Lazear MR, Forli S, Correia BE, Cravatt BF. Global profiling of lysine reactivity and ligandability in the human proteome. Nat Chem. 2017; doi:10.1038/nchem.2826

48. Ravichandran LV, Chen H, Li Y, Quon MJ. Phosphorylation of PTP1B at Ser50 by Akt Impairs Its Ability to Dephosphorylate the Insulin Receptor. Mol Endocrinol. 2001;15: 1768–1780. doi:10.1210/mend.15.10.0711

49. Alonso A, Sasin J, Bottini N, Friedberg I, Friedberg I, Osterman A, et al. Protein Tyrosine Phosphatases in the Human Genome. Cell. 2004;117: 699–711. doi:10.1016/j.cell.2004.05.018

50. Chen Y-NP, LaMarche MJ, Chan HM, Fekkes P, Garcia-Fortanet J, Acker MG, et al. Allosteric inhibition of SHP2 phosphatase inhibits cancers driven by receptor tyrosine kinases. Nature. 2016;535: 148–152. doi:10.1038/nature18621

51. Erlanson DA, McDowell RS, He MM, Randal M, Simmons RL, Kung J, et al. Discovery of a new phosphotyrosine mimetic for PTP1B using breakaway tethering. J Am Chem Soc. 2003;125: 5602–5603. doi:10.1021/ja034440c

52. Andersen HS, Olsen OH, Iversen LF, Sørensen ALP, Mortensen SB, Christensen MS, et al. Discovery and SAR of a novel selective and orally bioavailable nonpeptide classical competitive inhibitor class of protein-tyrosine phosphatase 1B. J Med Chem. 2002;45: 4443–4459. Available: https://www.ncbi.nlm.nih.gov/pubmed/12238924

53. Andersen HS, Iversen LF, Jeppesen CB, Branner S, Norris K, Rasmussen HB, et al. 2-(oxalylamino)-benzoic acid is a general, competitive inhibitor of protein-tyrosine phosphatases. J Biol Chem. 2000;275: 7101–7108. Available: https://www.ncbi.nlm.nih.gov/pubmed/10702277

54. Kabsch W, Wolfgang K. XDS. Acta Crystallogr D Biol Crystallogr. 2010;66: 125–132. doi:10.1107/s0907444909047337

55. Karplus PA, Diederichs K. Linking crystallographic model and data quality. Science. 2012;336: 1030–1033. doi:10.1126/science.1218231

56. Adams PD, Afonine PV, Bunkóczi G, Chen VB, Davis IW, Echols N, et al. PHENIX: a comprehensive Python-based system for macromolecular structure solution. Acta Crystallogr D Biol Crystallogr. 2010;66: 213–221. doi:10.1107/S0907444909052925

57. McCoy AJ, Grosse-Kunstleve RW, Adams PD, Winn MD, Storoni LC, Read RJ. Phaser crystallographic software. J Appl Crystallogr. 2007;40: 658–674. doi:10.1107/s0021889807021206

58. Chen VB, Arendall WB, Headd JJ, Keedy DA, Immormino RM, Kapral GJ, et al. MolProbity: all-atom structure validation for macromolecular crystallography. International Tables Online. 2012. pp. 694–701. doi:10.1107/97809553602060000884

59. Moriarty NW, Grosse-Kunstleve RW, Adams PD. electronic Ligand Builder and Optimization Workbench (eLBOW): a tool for ligand coordinate and restraint generation. Acta Crystallogr D Biol Crystallogr. 2009;65: 1074–1080. doi:10.1107/S0907444909029436

60. Long F, Nicholls RA, Emsley P, Graǽulis S, Merkys A, Vaitkus A, et al. AceDRG: a stereochemical description generator for ligands. Acta Crystallogr D Struct Biol. 2017;73: 112–122. doi:10.1107/S2059798317000067

61. Murshudov GN, Skubák P, Lebedev AA, Pannu NS, Steiner RA, Nicholls RA, et al. REFMAC5 for the refinement of macromolecular crystal structures. Acta Crystallogr D Biol Crystallogr. 2011;67: 355–367. doi:10.1107/S0907444911001314

62. Emsley P, Lohkamp B, Scott WG, Cowtan K. Features and development of Coot. Acta Crystallogr D Biol Crystallogr. 2010;66: 486–501. doi:10.1107/S0907444910007493

63. Schrödinger L. The PyMOL Molecular Graphics System, Version 1.8.

